# Large-scale pooled selection enables mapping of modifiers of RNAi gene knockdown effects in *C. elegans*

**DOI:** 10.64898/2026.10.08.757717

**Authors:** Stefan Zdraljevic, Alex Lee, Laura Walter-McNeill, Heriberto Marquez, Amy K. Webster, Rojin Chitrakar, Jameson Blount, L. Ryan Baugh, Joshua S. Bloom, Leonid Kruglyak

**Author notes:** Corresponding author: Stefan Zdraljevic, Joshua S. Bloom Leonid Kruglyak. Alex Lee: [ ], Laura Walter-McNeill: [ ], Heriberto Marquez: [ ], Amy K. Webster - [ ], Rojin Chitrakar: [ ], Jameson Blount - [ ], L. Ryan Baugh - [ ], Leonid Kruglyak: [ ].

## Abstract

Genetic interactions contribute to phenotypic variation, yet most studies characterize them in a small number of reference backgrounds rather than across natural populations. Studying genetic interactions in *Caenorhabditis elegans* is facilitated by RNAi by feeding, which enables gene-specific knockdowns across many genetic backgrounds without the need to construct individual mutant lines. However, individually screening hundreds of isolates across many RNAi conditions is laborious. Here, we developed a pooled platform that uses whole-genome sequencing to quantify the fitness of hundreds of genetically distinct *C. elegans* strains competing in a single flask. We leveraged this platform to evaluate RNAi responses in a pooled population of RNAi-sensitive *C. elegans* strains and identified two classes of QTL that influence RNAi responses. The first class influences responses to knockdowns of specific genes and is exemplified by a QTL that is specific to knockdown of *mig-6*. The second class influences general RNAi-response across multiple knockdowns. One QTL in this class localized to the right arm of chromosome III and was detected by both GWA and xQTL mapping, which suggests that a common large-effect variant underlies this QTL. We identified a T96K missense change in the dsRNA receptor SID-2 as the underlying causal variant. The 96K allele resides in the extracellular domain of SID-2 and confers increased RNAi sensitivity relative to the 96T allele. We hypothesize that the 96K allele enables greater uptake of ingested lumenal dsRNA through higher-affinity binding to the dsRNA phosphate backbone. The platform we developed enables systematic mapping of genetic interactions across natural populations at a previously impractical scale.

## Introduction

The same genetic lesion does not produce the same outcome in every individual who carries it. This shows up as incomplete penetrance and variable expressivity in Mendelian disease, where carriers of an identical pathogenic variant range from unaffected to severely affected. A growing body of work indicates that rare and common background variation is a major source of this variation [1,2]. For example, Huntington’s disease risk is conferred entirely by an *HTT* CAG repeat expansion, but background variation with two independent effects on chromosome 15 moves onset 6.1 years earlier and 1.4 years later, and a chromosome 8 locus hastens onset by 1.6 years [3]. Biobank-scale sequencing is beginning to identify rare large-effect modifiers, but such modifiers remain difficult to detect because isolating the effect of one variant against a fixed, inherited background requires exceedingly large samples [2]. These observational studies establish that genetic background is a major determinant of disease phenotypes, but they remain observational because neither the causal mutation nor background can be chosen or resampled.

Model organisms don’t have these constraints because a single perturbation can be introduced into many natural genetic backgrounds and its effect measured directly. Modifier screens at scale show that dozens of essential genes in the laboratory reference yeast strain are dispensable in at least one natural genetic background [4,5]. These studies also reveal that the genetic modifiers of knockout mutations tend to map to single modifier genes. However, extending these studies to metazoan organisms is largely intractable because it requires constructing hundreds of individual mutant lines. *Caenorhabditis elegans* doesn’t have this limitation because RNAi by feeding enables knock down of target genes without altering the genome [6,7]. Despite the unusual sensitivity of *C. elegans* to ingested RNAi [8], individually measuring hundreds of strains across many knockdown conditions has limited the breadth of this approach. The most extensive such screen assayed 29 maternal-effect genes across 55 wild strains and recovered highly polygenic architecture driven by gene-specific modifiers [9].

Pooled selection and sequencing can enable simultaneous phenotyping of many genotypes in a single flask. In *C. elegans*, this has been implemented using molecular inversion probes (MIP-seq) to track strain identity within a pooled competition. This approach was implemented to study variation in competitive fitness of 22 *C. elegans* isolates in the presence of microsporidia [10], and resolved a starvation-resistance locus identified by pooled phenotyping of 100 genetically diverse strains [11]. A major limitation to this approach is the construction of population-specific probe panels before phenotyping can begin, which can limit panel size and ties reagents to one fixed set of strains. By contrast, inferring strain frequencies directly from whole-genome sequencing only requires sequenced genotypes that are publicly available on CaeNDR [12,13].

Extending pooled selection experiments to identify genetic modifiers of a specific knockdown must contend with heritable variation in RNAi sensitivity across the *C. elegans* population [9,14–16]. A panel of wild isolates therefore segregates general RNAi modifiers that shift responses of every knockdown and knockdown-specific modifiers. We therefore designed an experiment that could simultaneously map general RNAi-sensitivity variation and identify potential genetic modifiers of knockdown perturbations.

Here we present a pooled phenotyping platform for *C. elegans* that infers strain frequencies from bulk sequencing using non-negative least squares regression and show that it performs comparably to a previously published MIP-seq experiment [11]. We used this platform to phenotype 231 wild *C. elegans* isotypes exposed to double-stranded RNA targeting *pos-1* and identified multiple QTL that explain RNAi-response variation across the population. Of the identified QTL, one that localizes to the right arm of chromosome III was independently identified in an xQTL experiment [17,18]. We fine-mapped this region to a single residue substitution, T96K, in the extracellular domain of the dsRNA receptor SID-2. We show that this variant increases RNAi sensitivity in multiple genetic backgrounds, including the standard laboratory strain, N2. The T96K shifts responses to multiple knockdowns because it likely affects RNAi delivery to the intestine. Contrasting two knockdown conditions isolates the target-specific component and reveals multiple large-effect loci modifying the response to a single gene knockdown.

## Results

### Accurate strain frequency inference from whole-genome sequencing

To enable selection screens of large genetically diverse populations, we developed a method that uses whole-genome sequencing to accurately infer individual strain frequencies from a pooled set of isolates with known genotypes. We first tested this method by simulating strain phenotypes and genotype calls for a range of sequencing depths (1-500x). We observed that we could accurately recover strain frequencies using a regression-based approach (Methods) [19], with close agreement between input and recovered strain frequencies (median *r²* of 0.906 at 10X depth and 0.986 at 30X depth; **Figure S1.**) We applied this method to real pooled populations and found that we were able to accurately recover input frequencies (**Figure S2;** Pearson *r²* = 0.994). Finally, we tested the method against a published pooled-phenotyping experiment that used molecular inversion probes to infer strain frequencies of 100 pooled *C. elegans* stains [11]. We performed whole-genome sequencing on the published samples and inferred strain frequencies using our method. We found good agreement in frequency estimates between the methods across 23 individual samples (average Spearman’s ρ = 0.83, **Figure S3A-B**), and a composite trait that represents strain recovery post L1 starvation (Spearman’s ρ = 0.97) (**Figure 1A**). Taken together, these three validation experiments demonstrate that the method can accurately infer strain frequency dynamics and enable pooled selection without strain-specific barcodes or probes.

**Figure 1:**
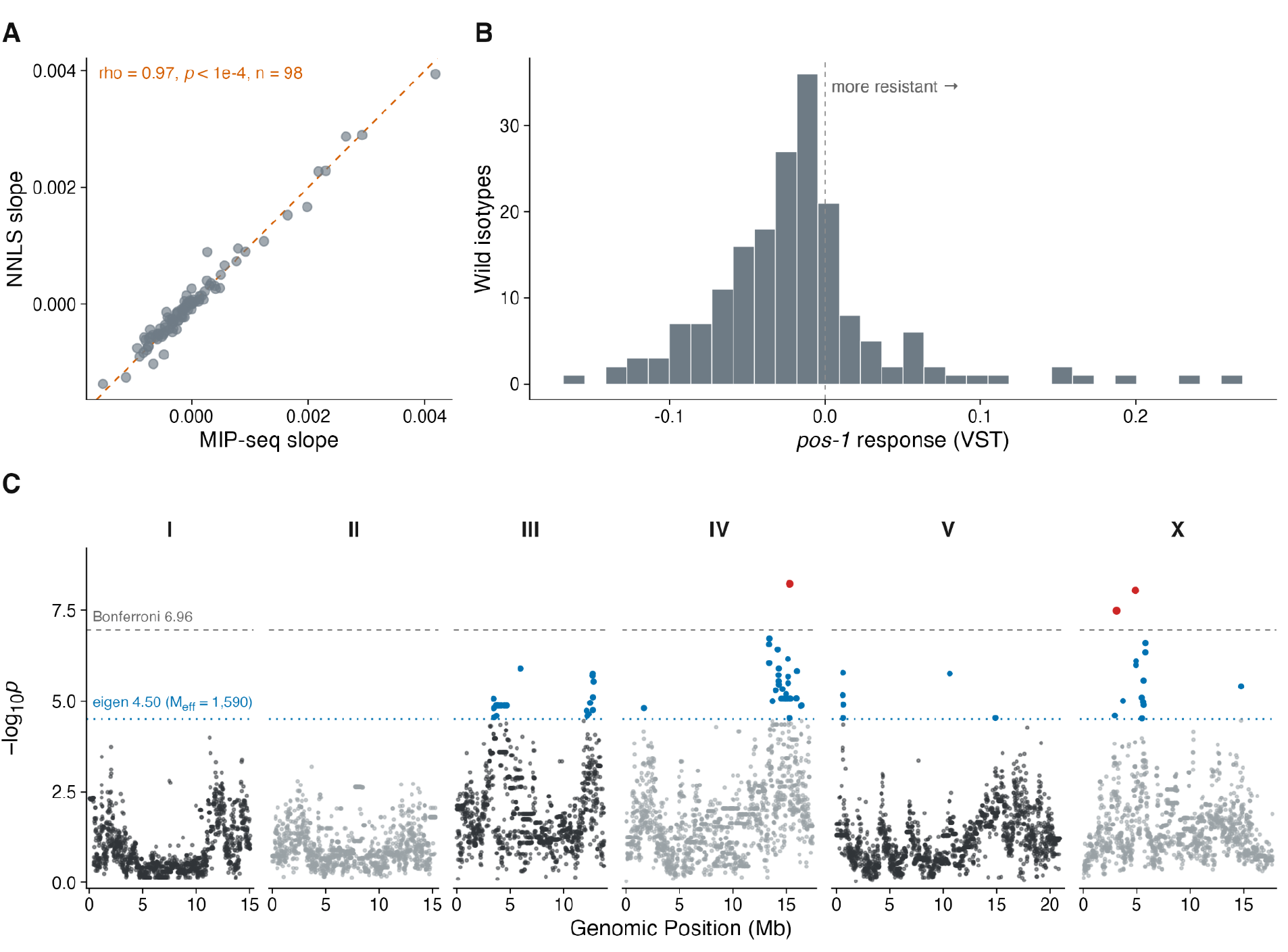
Pooled phenotyping maps multiple RNAi-sensitivity loci. A) Comparison of MIP-seq (x-axis) and NNLS (y-axis) per-strain rate of change in pool frequency after L1 starvation. Each dot represents a strain’s rate of change after starvation, averaged across five replicates. The dashed line is y=x. B) Distribution of the pooled *pos-1* RNAi responses across wild isotypes. The dashed line marks the zero; positive values represent *pos-1* resistant strains that increase after *pos-1* treatment. C) Genome-wide association scan for the phenotype displayed in B. Each dot represents an SNV. The y-axis represents the strength of association (-log10*p*) for each SNV distributed along the genome (x-axis). The dashed lines represent thresholds for multiple testing correction (blue = eigen-decomposition; grey = Bonferroni).

### Pooled phenotyping enables evaluation of wild isolate RNAi responses

*C. elegans* RNAi screens are routinely performed in the N2 strain. Extending these screens to multiple wild isolates is challenging because each isolate needs to get assayed independently. We reasoned that pooled phenotyping would enable fast evaluation of wild isolate RNAi responses. N2 strains exposed to *pos-1* RNAi exhibit an embryonic lethal phenotype[20], making this treatment amenable to the pooled phenotyping because RNAi-responsive strains won’t contribute progeny to future generations. To test if we could easily evaluate wild isolate RNAi responses with pooled sequencing, we performed a pilot experiment in which we exposed a pool of 224 wild isolates to *pos-1* or control bacteria for two generations across four replicates (Methods). We found good agreement between strain frequencies across the replicate conditions (**Figure S4**; Spearman’s ρ = 0.83–0.90).

Of the 224 isolates in the pool, we detected 184 individuals in both control conditions. We found that 76% of these isolates declined in frequency in *pos-1* RNAi relative to the control condition, more than would be expected if frequency changes reflected noise alone (binomial test, p = 7.3 × 10⁻¹³) (**Figure 1B**). Because pool frequencies are compositional, this proportion measures decline relative to the pool, rather than absolute per-strain responses. In total, 53 of these 184 isolates are absent from every *pos-1* pool while present in both controls. Of the remaining 131 isolates, 52 declined in frequency (one sample t-test on four replicate deltas q < 0.05), showing that 105 of 184 isolates (57%) are at least partially responsive to *pos-1* RNAi. We performed a genome-wide association scan on the *pos-1* responses of these 184 isolates, which identified two QTL: one on the right arm of chromosome IV and one on the left arm of chromosome X (**Figure 1C**). This analysis does not determine whether these QTL are *pos-1*-specific or general RNAi-response QTL. We reasoned that if they are general RNAi-response QTL, we would want to minimize their contribution to future experiments.

To enable subsequent screens, we sought to construct a pooled population of RNAi-responsive strains. We manually re-evaluated *pos-1* RNAi responses of 191 wild strains on agar plates and identified 93 strains with robust RNAi responses that we included in the pool (Methods).

### Knockdown-specific and general RNAi response QTL segregate in wild isolates

Having assembled a pool of RNAi-responsive strains, we next asked whether we could identify QTL that modify responses to individual RNAi treatments. We grew this pool on bacteria expressing double-stranded RNA against *pos-1* and *mig-6* and on the HT115 empty-vector control, sequenced the resulting populations, and inferred strain frequencies (Methods). We performed genome-wide association scans on the responses to these knockdowns and identified QTL for *pos-1* and *mig-6* RNAi (Figure 2). The *pos-1* response mapped to the right arm of chromosome III (III:12,353,680, -log10p = 7.04, allele frequency 0.11), just 0.4 Mb from the association we identified in our pilot experiment (**Figure 1C**). The *mig-6* response mapped to the center of chromosome V (V:14,647,434, -log10p = 7.71, allele frequency 0.13), a locus we did not identify in either *pos-1* experiment, suggesting that this locus is *mig-6*-specific.

**Figure 2:**
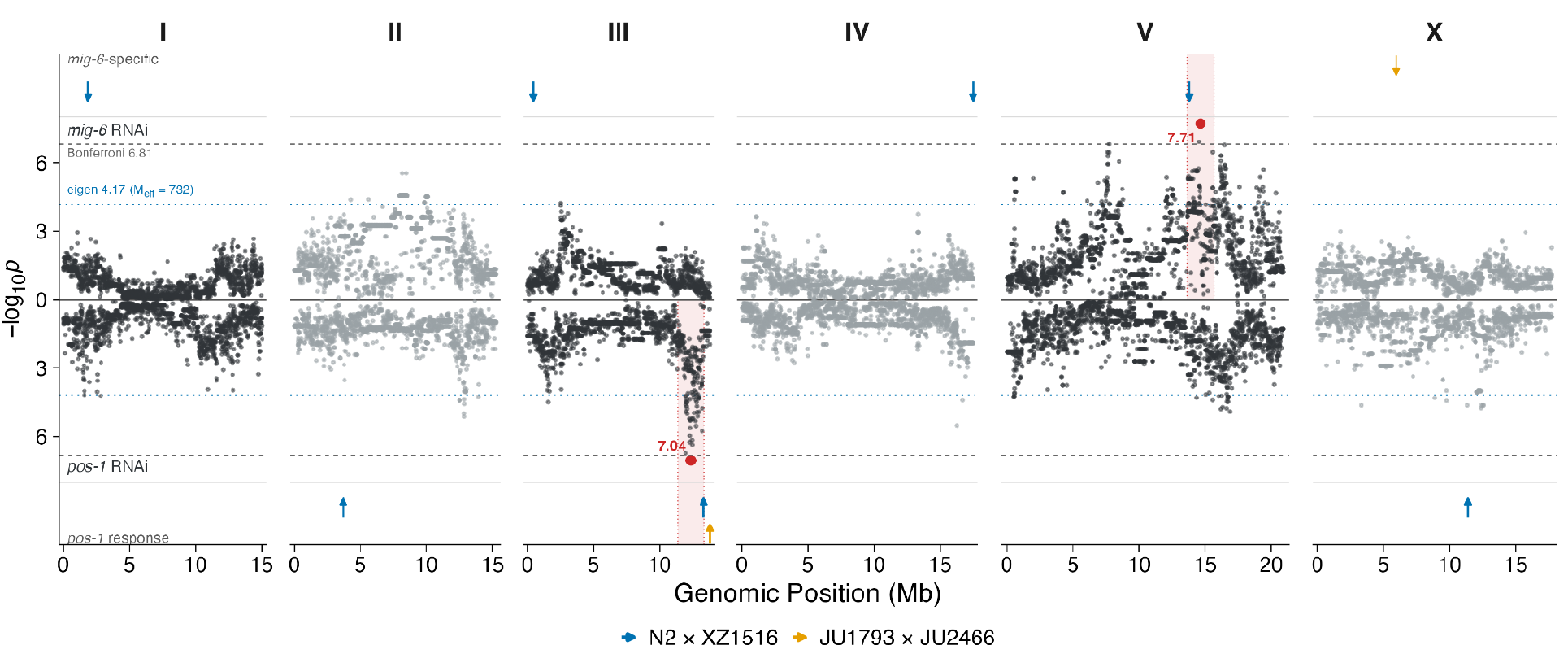
An RNAi-sensitive population identifies knockdown-specific modifier QTL. Genome-wide association scan for strain responses to *mig-6* (top) and *pos-1* (bottom) RNAi. Each dot represents an SNV. The y-axis represents the strength of association (-log10*p*) for each SNV distributed along the genome (x-axis). The dashed lines represent thresholds for multiple testing correction (blue = eigen-decomposition; grey = Bonferroni). Arrows at the top and bottom of the plot indicate the position of *pos-1*- and *mig-6*-specific QTL identified by xQTL. Blue arrows from the N2 x XZ1516 cross and orange arrows from the JU1793 x JU2466 cross.

We next asked if additional genetic variation contributed to exceptionally sensitive and resistant strains in these pooled knockdown experiments. Evaluating knockdown responses of recombinant progeny from phenotypically divergent parents could reveal variants the pooled scan missed, so we identified strains to construct large cross populations for bulk segregant mapping (Methods). One cross was constructed between JU1793, which was resistant to both *mig-6* and *pos-1* RNAi, and JU2466, which was sensitive to *pos-1* RNAi and had an average *mig-6* response (**Figure S5**). We made use of a previously constructed cross between N2 and XZ1516 because XZ1516 had among the strongest responses to both RNAi conditions (**Figure S5**) [18]. Each cross was continued for 10 generations and then exposed to RNAi. We sequenced the cross populations after two generations on the RNAi conditions and used allele frequency deviations between conditions to perform QTL mapping. By comparing parental frequencies after RNAi exposure to the control HT115 condition, we identified multiple QTL shared between the two crosses, as well as cross-specific QTL (**Figure S6B-C**, **Table S14**). Some QTL were shared between the two RNAi conditions, and we refer to them as general RNAi-response QTL. By contrasting allele frequencies after *pos-1* and *mig-6* RNAi exposure, we identified gene-specific QTL.

One major general RNAi-response QTL we identified in both crosses localized to the right arm of chromosome III (**Figure S6B-C**), in close proximity to the QTL we observed in the pooled association mapping (**Figure 2**). We also identified a large *mig-6*-specific QTL in the XZ1516 cross that co-localized with the *mig-6*-specific QTL we identified in the pooled experiment. In addition to these re-discovered QTL, we identified two novel large-effect *mig-6*-specific QTL: one on chromosome I in the N2 x XZ1516 cross, where parental allele frequency differs by 0.391 (95% CI 0.378-0.404) between the *mig-6* and *pos-1* conditions, and one on chromosome X in the JU1793 x JU2466 cross, with a parental allele frequency difference of 0.713 (95% CI 0.679-0.747). We conclude that these loci are *mig-6* knockdown-specific because their parental allele frequencies only deviate after *mig-6* exposure. Taken together, these results highlight the power of the pooled-phenotyping approach to measure RNAi responses across the C. elegans population, point to substantial RNAi-response variation segregating in the population, and map several loci that modify gene-specific knockdown responses.

### A high-frequency variant in the dsRNA transporter SID-2 confers increased sensitivity to ingested RNAi

We next sought to uncover the variant underlying the general RNAi-response QTL on the right arm of chromosome III. We first confirmed that JU2466 had distinct *pos-1* RNAi phenotypes using a plate-based embryonic lethality assay (p = 1.4 × 10⁻⁴⁹) (**Figure S7**; Methods). Next, we introgressed the chromosome III QTL region from JU2466 into the JU1793 genetic background and found that it conferred *pos-1* sensitivity to JU1793 at a level that was not significantly different from that of JU2466 (p = 0.49)). Through iterative near-isogenic line (NIL) construction, we localized the QTL to a 37 kb interval spanning 13.658–13.695 Mb on chromosome III (**Figure 3C**; **Figure S7**). Within this region, we found 27 sequence differences in twelve genes between the parent strains. Only two of these differences altered a protein sequence, and both were missense mutations in *sid-2* (V5L and T96K). *sid-2* encodes a single-pass transmembrane protein localized to the apical membrane of intestinal cells that facilitates uptake of dsRNA from the gut lumen, making it an excellent candidate to harbor variation that contributes to RNAi sensitivity across wild isolates. We focused on the T96K variant for two reasons. First, it is also the variant that distinguishes the parents of the N2 x XZ1516 cross, in which an overlapping general RNAi-response QTL is segregating that we hypothesized may be driven by the same underlying variant (**Figure S6B-C**). Second, residue 96 is predicted to reside in the extracellular domain of SID-2 and has the potential to alter interactions with double-stranded RNA.

**Figure 3:**
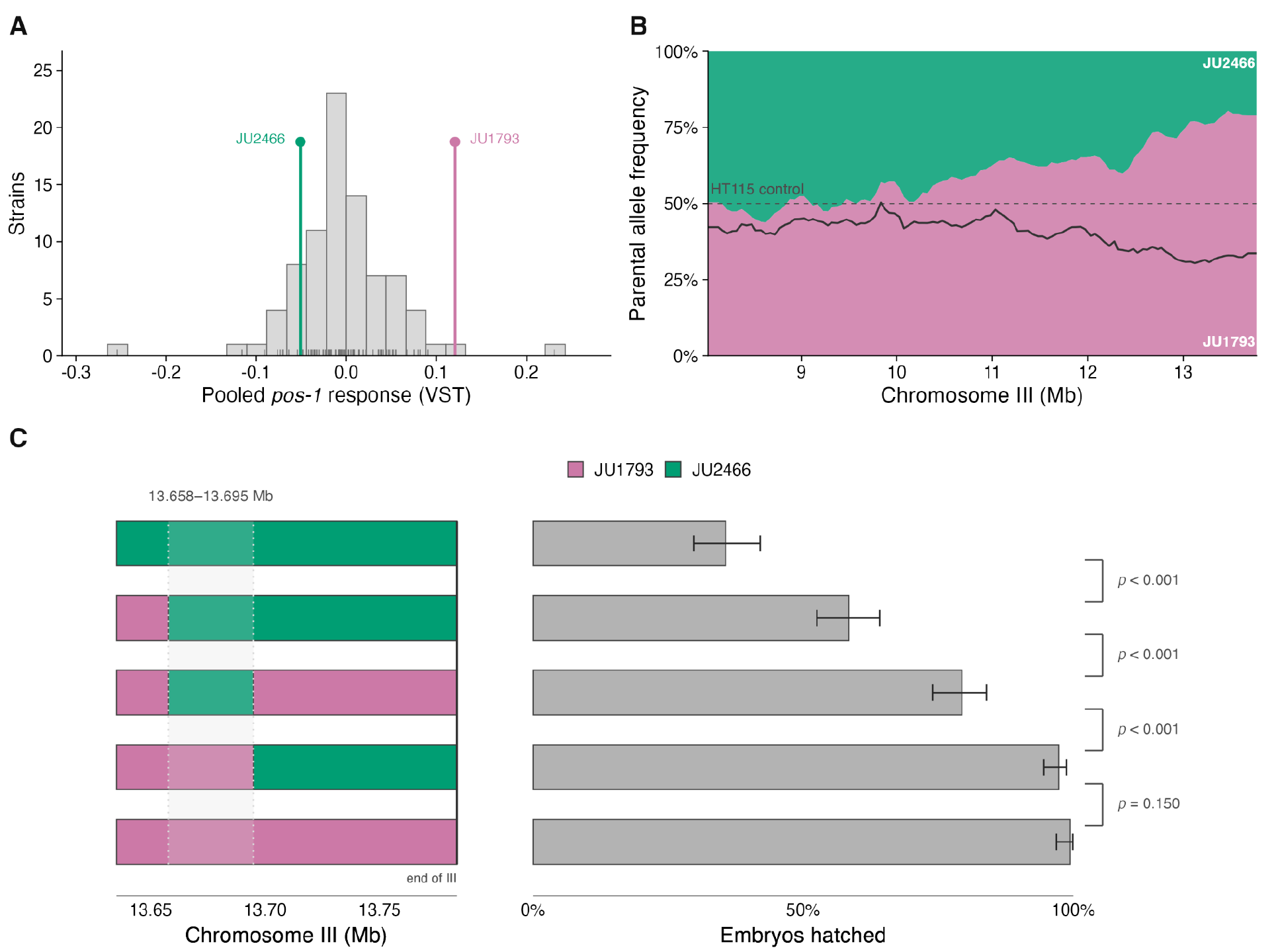
NILs resolve an RNAi-sensitivity QTL. A) *pos-1* RNAi responses from the pooled phenotyping of the RNAi-sensitive population. JU2466 and JU1793 are highlighted in green and pink, respectively. B) The fraction of each JU2466 and JU1793 allele on the y-axis across the right arm of chromosome III (x-axis) from the JU2466 x JU1793 crosses after two generations on *pos-1* RNAi. The black line represents the parental allele fraction in the empty vector control. C) NIL genotypes across the chromosome III RNAi-sensitivity QTL are shown on the left. *pos-1* RNAi responses expressed as percent of embryos hatched are shown on the right, where fewer embryos hatched corresponds to more RNAi sensitivity. Bars are Wilson 95% binomial intervals. Brackets in the right-hand column are Fisher’s exact tests on the hatched/unhatched counts of adjacent rows, Holm-corrected over the four comparisons,

We generated reciprocal allele-swap strains to test whether this variant contributed to *pos-1* RNAi responses. Introducing the 96K allele into the JU1793 strain lowered its hatching rate from by 41.7%, (p = 1.7 × 10⁻²⁴), while introducing the 96T allele into the JU2466 strain raised its hatching rate by 13.9% (p = 3.3 × 10⁻⁸)(**Figure 4A**). Therefore, the 96T reference allele produces a weaker RNAi response than the alternate 96K allele in both parental backgrounds. Because N2 is the standard laboratory reference strain and has been used in countless successful RNAi experiments, we were surprised to find that the 96T reference allele caused a weaker RNAi response. We therefore asked if we could increase the RNAi sensitivity of N2 by introducing the 96K allele. Under our standard assay conditions (50:50 mixture of *pos-1* and HT115 bacteria), introducing 96K into the N2 strain produced no difference in the embryo hatching rate. This is likely because at this dose, N2 fails to hatch 94% of its embryos (6% hatched, 95% CI 3.2-10.0), leaving little room for further sensitization (**Figure S8**). When we lowered the *pos-1* bacteria concentration on the assay plates, we found that the 96K allele lowered the N2 hatching rate from 32.3% (95% CI 26.4-38.7) to 4.0% (95% CI 2.6-6.0) (two independently edited lines, 4.4% and 3.7%; **Figure 4B**). Thus, this allele also increases RNAi sensitivity in this genetic background.

**Figure 4:**
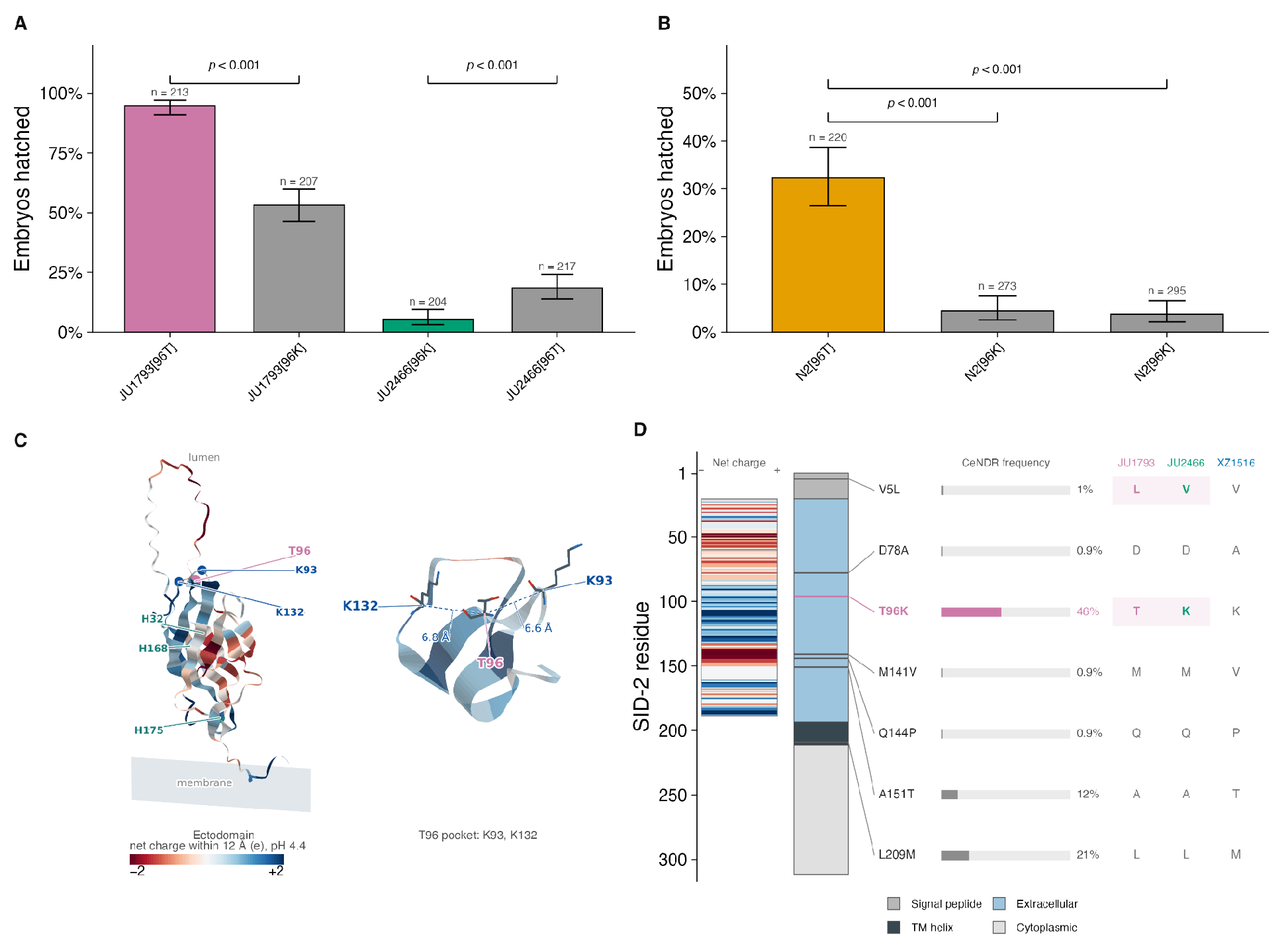
The SID-2[T96K] allele underlies variable *C. elegans* RNAi responses. A) The fraction of embryos hatched after *pos-1* RNAi exposure is shown on the y-axis for JU1793 (pink), JU2466 (green), and the two allele swap strains in gray (JU1793[96K] and JU2466[96T]). Fisher’s exact test on the counts, each edit against the unedited allele in the same background: JU1793 *p* = 1.7e-24 and JU2466 *p* = 3.9e-05. Bars are Wilson 95% binomial intervals. B) The fraction of hatched embryos after *pos-1* RNAi is shown for N2 and N2 with the 96K allele is shown. Fisher p = 7.8e-17 and p = 6.2e-19. C) The SID-2 ectodomain coloured by side-chain charge of all ectodomain residues with a C-alpha within 12 Å at the gut-lumen pH of 4.4, red negative and blue positive. Left, residues 21-188 of chain A, showing T96 (pink, the JU1793 allele the model carries), the two lysines that make its pocket, K93 and K132 (blue), and the three extracellular histidines H32, H168 and H175 (teal) known to affect dsRNA uptake. Right, the pocket enlarged, with C-alpha distances from T96 to K93 (6.6 A) and K132 (6.8 A). D) *sid-2* coding variation across the wild population that differentiates JU1793, JU2466, and XZ1516. The protein runs vertically with residue 1 at the top and is colored by the predicted domain. Net charge from C is shown to the left. Each protein-altering variant is marked to the right with population-wide allele frequency, followed by three columns giving the allele state in the three strains included in crosses.

96T is the third position of a predicted N-linked glycosylation sequon (consensus N-x-S/T), N94-C95-T96. The 96K allele destroys the sequon and adds a positive charge. Because N-linked glycosylation of membrane proteins typically supports folding, stability, and membrane localization, and because this sequon is conserved across nine of ten species of the *elegans* supergroup and its sister *C. kamaaina* (**Figure S9A**), loss of the site would be predicted to impair SID-2 function. Although there is no experimental evidence of N94 being glycosylated in *C. elegans*, the effect we observe goes in the opposite direction, with the 96K allele increasing sensitivity to ingested double-stranded RNA relative to 96T.

To separate the loss of the sequon from the introduction of the lysine, we introduced a N94A substitution into both JU1793 and JU2466. The N94A change caused a decrease in *pos-1* RNAi responses in both genetic backgrounds, consistent with disrupted SID-2 function (**Figure S9B**). Because both alleles break the sequon in JU1793, these results indicate that the charge introduced by the lysine reverses the effect of losing the glycosylation sequon. Since the sequon is already disrupted by the 96K allele in JU2466, this observation suggests that the asparagine contributes to SID-2 function independently of an attached glycan. Consistent with this hypothesis, the N94A substitution in the JU2466 background conferred greater RNAi resistance than did restoring the ancestral residue at position 96 (42.5% versus 18.4%, OR 3.3, p = 9e-8). This result is difficult to reconcile with the two residues acting through a shared modification.

We hypothesized that by introducing a positively charged residue, the 96K allele might facilitate a stronger interaction with the phosphate backbone of double-stranded RNA and therefore better uptake of double-stranded RNA. To assess this, we computed the local net charge in 12 Å windows across the SID-2 ectodomain. The window centered on residue 96 has a net charge of +1.24e. This is the 82nd percentile of all ectodomain residues and is largely driven by K93 and K132, which sit 6.6 and 6.8 Å away. (**Figure 4C**, **Figure S10**). Introducing the 96K variant to this local region shifts the net charge to +2.24 e, which corresponds to the 98th percentile of the ectodomain’s local net charge distribution (**Figure S10**). While these observations don’t prove that the 96K allele is causing a stronger interaction with double-stranded RNA by increasing the net charge of its local environment, it does agree with previous reports that modifying critical histidines in the ectodomain to positive arginines is less disruptive to uptake of double-stranded RNA than modifying the histidines to alanines [21].

The 96K allele is found in 46.3% of sequenced *C. elegans* strains and is always found with the 153T allele (**Figure 4D, Figure S11**). JU1793 is one of eleven strains that carry the 96T and 153T alleles. Because JU1793 and JU2466 differ only at T96K and V5L, the cross likely captured all of the RNAi-response variation attributable to SID-2 coding differences between them (**Figure S6**). XZ1516, however, carries five additional missense variants in SID-2, four of them in the extracellular domain (**Figure 4D**), which might further modify dsRNA uptake. Across the 684 isotypes in CaeNDR, we found 20 protein-altering variants at 19 sites in SID-2, including a frameshift in a single isotype, ECA2199, that truncates the protein before its transmembrane domain (**Figure S11)**. This variation may contribute to further natural differences in responses to ingested RNAi.

## Discussion

We developed a platform that can reliably infer the frequencies of hundreds of strains in pooled populations. This platform can readily be applied to any pooled population of strains with known genotypes because it relies on whole-genome sequencing. We first used this platform to map RNAi-response variation across 231 *C. elegans* isolates and identify three distinct QTL (**Figure 1B-C**). We then applied it to an 84-isolate pool of RNAi-sensitive individuals that identified the same general RNAi-response QTL on chromosome III, as well as an additional gene-specific QTL on chromosome V (**Figure 2**). We validated these QTL using two independent bi-parental crosses that revealed several other modifiers of *mig-6* and *pos-1* RNAi knockdowns (**Figure S8B-C**).

We resolved the general RNAi-sensitivity locus on the right arm of chromosome III to a T96K missense mutation in the extracellular domain of the dsRNA receptor SID-2. We showed that strains with the 96K allele are more sensitive to *pos-1* RNAi than strains with the 96T allele across three genetic backgrounds, including the standard laboratory strain N2 (**Figure 4A-B**). T96 sits in a conserved N-glycosylation sequon, N94-C95-T96, and while the 96K allele adds a positive charge to the ectodomain that might facilitate stronger interaction with the phosphate backbone of double-stranded RNA, it also disrupts the conserved sequon. Because N-glycosylation is associated with protein stability and localization, disrupting the sequon is expected to impair SID-2 function. However, we observed the opposite effect in the T96K allele swap experiments. We therefore introduced a N94A substitution in both genetic backgrounds to disrupt the sequon independently of T96K. The 94A allele makes JU1793 and JU2466 more resistant to *pos-1* RNAi, suggesting that this allele impairs SID-2 function (**Figure S10**). Because T96K also disrupts the sequon while contributing a positive charge, we hypothesize that the electrostatic effect of T96K must be large enough to override the antagonistic, glycosylation-linked resistance effect we observed in the N94A edited strains. By adding localized positive charge to the ectodomain, the lysine allele may facilitate more double-stranded RNA uptake than the threonine, thereby inducing a stronger RNAi response in carrier worms. While a previous study did not detect glycosylation of the N94-C95-T96 sequon[22], this lack of detection is likely driven by sampling depth in the proteomics dataset in which only 870 proteins were recovered [22]. In proteomics studies that do detect SID-2, only the cytoplasmic domain is detected, not the N94-C95-T96 peptide [23,24].

Sensitivity to ingested dsRNA has been gained and lost repeatedly across *Caenorhabditis* [8], which suggests that the trait is under selection and may explain why we observe a high-frequency large-effect variant in the receptor responsible for internalizing dsRNA from the environment. A source of selective pressure may be the RNA viruses that individuals encounter in the wild, because *C. elegans* antiviral defense is mediated by RNAi [25]. Natural variation in *C. elegans* antiviral responses is already known to be consequential: a deletion in *drh-1*, the worm RIG-I homolog, disables viral RNA dicing and antiviral immunity in some wild isolates [26]. Another source of selective pressure may be small RNAs from bacteria, which have been reported to influence *C. elegans* behavior [27]. Different versions of the receptor that imports extracellular RNA across the gut could therefore be favored depending on the composition of the local microbial community.

In addition to the identification of general RNAi-response QTL, the pooled screen and a bi-parental cross revealed two *mig-6*-specific QTL localized to the center of chromosome V and the left arm of chromosome I (**Figure 2**, **Figure S8**). We hypothesize that these QTL are *mig-6*-specific because both N2 and XZ1516 are strong responders to RNAi, making disrupted RNA transport to *mig-6*-expressing somatic tissues a less-likely explanation. The chromosome V QTL is likely a common variant of large effect because it was identified in the pooled screen, while the variant underlying the chromosome I QTL is likely less frequent across the mapping population because it was only detected in one cross and not in the pooled screen. Establishing that these QTL are *mig-6*-specific would require identifying the underlying variants or performing additional RNAi-knockdown experiments in this cross to verify the lack of a chromosome I and V signal in unrelated RNAi conditions.

The mapping results highlight the ability of our platform to identify large-effect genetic modifiers of genetic perturbations. Thousands of rare Mendelian diseases are caused by loss-of-function variants, and their penetrance and expressivity are frequently modified by background genetic variation [2]. Such modifiers are currently difficult to identify in human genetics studies because rare disease-causing variants are carried by too few individuals to give modifier scans sufficient statistical power. The platform we developed enables us to identify genetic modifiers for a wide range of perturbations of worm homologs of disease-causing genes in pools of genetically diverse *C. elegans* isolates.

## Methods

### Strain Maintenance

Wild isolates were obtained from The Caenorhabditis

Natural Diversity Resource (CaeNDR) [12,13]. Prior to plating on RNAi conditions, all *C. elegans* strains were maintained at 20°C on modified nematode growth medium (NGMA), containing 1% agar and 0.7% agarose and seeded with OP50 [29].

### RNAi Bacteria

All RNAi bacteria were streaked onto LB + 100 µg/ml carbenicillin agar plates to form single colonies. Single colonies were grown overnight in 3 ml liquid LB + 100 µg/ml carbenicillin cultures. 24 hours later, the cultures were diluted 1:1000 in LB + 100 µg/ml carbenicillin and grown for 16 hours. Finally, RNA expression was induced with 1 mM IPTG for four hours. 100 µl of these cultures were used to seed 6 cm plates and 1000 µl for 10 cm plates. RNAi plates were dried for two days at room temperature and then transferred to 4°C [30,31].

After observing inconsistent RNAi responses in known responding strains, large volumes of RNAi bacterial cultures were grown, OD-standardized, and stored at 4°C in aliquots to reduce batch variability. Three liters of each culture were transferred to centrifuge bottles and centrifuged at maximum speed for 10 minutes. The supernatant was discarded, and the remaining bacterial pellet weight was calculated by subtracting the empty bottle weight from the pellet-containing bottle weight. The bacterial pellets were resuspended with K-media [32] at a ratio of 1 g : 4 mL to generate 10x RNAi bacterial stocks [33]. For plating convenience, stocks were further diluted to 1x and frozen at -80°C in 5 mL aliquots. All subsequent NIL phenotyping has been plated on NGMA + carbenicillin plates with a 1:1 mixture of *pos-1* and HT115 cultures normalized at an OD_600_ of 12.9.

### NNLS Simulations

Genotypes for *C. elegans* wild isolates were obtained from the CeNDR release (2021-01-21) in PLINK [34] binary format and read into R with the BEDMatrix package. Variants on each of the six chromosomes (I–V, X) were loaded separately and combined into a single genotype matrix G of dimension *m* markers × *n* strains. Missing calls were set to the reference state, and genotypes were binarized (homozygous-alternate coded as 1, reference as 0), which is appropriate given that wild isolates are nearly fully homozygous. From this matrix we precomputed the strain-by-strain Gram matrix **GᵀG** for use in the deconvolution step. Strain frequencies were estimated from pooled sequence data by non-negative least squares (NNLS) using RcppML::nnls.

Each wild isolate was assigned a fitness from one of seven published *C. elegans* traits with validated QTL (Albendazole_q75.TOF, PC1, assay_norm, mtDNA_ratio, value, amsacrine_f.L1, etoposide_median.TOF)[35–38], giving seven simulated populations of 327 strains. The trait value was the fitness, with a constant added to each trait such that fitness is always non-negative; a strain with no value for a trait was assigned fitness zero and is absent from that trait’s pool. Expected pooled allele frequencies were computed from the CeNDR genotype matrix, observed alt-allele counts were simulated by binomial sampling at 1, 3, 5, 10, 30, 50, 100 and 500x, and those counts were deconvolved and compared with the known input. Recovery was the squared Pearson correlation (r²) between estimated and true frequencies, computed over the strains present in each pool. The simulation was repeated ten times with independent binomial draws (seed 20260909); Figure S1 reports the mean r² across replicates with a band spanning the replicate range.

### Creation of pooled populations

We created pooled populations of approximately 48 strains each. To create these pooled populations, we recovered individual CeNDR strains on 6cm NGMA plates for three days and then bleached them on new 6cm plates. The following day, L1s were transferred to new 6cm plates and were allowed to starve. Once all strains from a given plate were starved (∼4 days), approximately 1 cm chunks from the starved plates were transferred to three different 10cm plates and grown for three days. After three days of growth, the worms from all the plates were washed off with M9 and put in a 50ml conical tube. Worms were then bleached to harvest embryos, and the embryos were resuspended in K media and arrested overnight as L1s. L1s were titered and diluted to contain approximately 1 million worms/ml and frozen for long-term storage. In total, we prepared eight of these populations, which we labeled A-H.

### Dilution experiment

To perform the dilution experiment, we recovered the B and C pooled populations from the freezer in 1 L flasks that contain 200ml of K media and 1X HB101. We allowed the populations to grow for 72 hours until they were gravid, at which point we gravity settled the worms in 50 ml conical tubes and bleached the population. We arrested the populations in K media overnight and allocated approximately 500,000 worms for a DNA preparation. We combined the two DNA preparations at nine known ratios (B1 fraction: BC1 = 0.09, BC2 = 0.17, BC3 = 0.29, BC4 = 0.44, BC5 = 0.62, BC6 = 0.76, BC7 = 0.86) and prepared libraries for whole-genome sequencing.

### Inferring strain frequencies

Raw sequencing reads were aligned to the N2 genome (build: WS276) using *bwa mem* [39]. The GATK *ASEReadCounter* tool was run on the resulting BAM files using the CaeNDR VCF (release: 20220216) [12]. For a given experiment, a given variant site was retained if it had a mean depth between 5× and 100× across all samples and no missing depth values. Next, we removed nonvariable sites in the experimental strain population and sites with fewer than 1,000 genotype calls in the VCF to prevent inclusion of sites without consistent calling across the population. To correct for reference bias in allele frequency estimation, variants with a minor allele frequency greater than 50% in the experimental population were recoded so that the more common allele was always treated as the reference, with corresponding allele counts flipped accordingly. Strain frequencies in each bulk sample were estimated by non-negative least squares (NNLS) using RcppML. The genotype matrix (markers × strains) and per-sample alternate allele counts were used to solve for strain composition by minimizing the residual between observed allele counts and those predicted by the linear combination of strain genotypes, with solutions constrained to be non-negative and normalized to sum to one. Estimated strain frequencies were output for all samples as a strains × samples matrix.

### MIP-seq comparison

We assessed the accuracy of NNLS deconvolution against published molecular inversion probe sequencing (MIP-seq) frequencies for the L1 starvation time course [11], across the 23 samples in which both measurements exist. The authors excluded CX11262, ECA348, and NIC260 from their analysis and so did we. One isolate included in the pool, PB306, is not a reference isotype in the CeNDR VCF, so we excluded it from the analysis as well. Finally, we excluded N2 because it is the reference strain that genotypes are called against and contains no ALT sites in the VCF. This brings the total of analyzed strains to 98. For each strain and replicate we subtracted the day 1 frequency from the frequency on each day and regressed that difference on day. Day 17 was excluded. Slopes were calculated for each replicate and then averaged, to give a “difference-based slope”. The identical calculation was applied to the pooled-sequencing and MIP-seq frequencies, so the two slopes estimates only differ in their input frequencies. Agreement was quantified by Spearman correlation across strains (ρ = 0.974, n = 98, p < 1e-4). Uncertainty was estimated by resampling markers with replacement across 100 bootstrap replicates of the deconvolution.

We also compared traits derived from our frequencies with the published phenotypes. Following Webster et al., we computed log2(f_day / f_baseline) for each strain and sample.. Slope is the regression of this ratio on day with Day 17 excluded; PC1 is the first principal component of the same ratios with day 17 included (prcomp, scaled and centered). Because the deconvolution returns exact zeros where MIP-seq read ratios do not, frequencies were floored at 1/(4n) before taking logarithms. The resulting traits agree with the published Slope at ρ = 0.890 and with the published PC1 at ρ = 0.822. The difference-based slope used above, agrees with the published log2(f_day / f_baseline) Slope at ρ = 0.888. The comparisons are detailed in Supplementary Note 1.

Sequencing depth requirements were assessed by randomly subsampling, without replacement, the reference and alternative allele observations at each marker to a target depth, rerunning the deconvolution, and comparing to the full-depth frequencies of the published traits. Target depths were 0.25, 0.5, 1, 3, 5 and 10x.

### Pooled GWA

#### Association mapping

Association mapping used GEMMA 0.98.5 with leave-one-chromosome-out (LOCO) kinship, run as a Nextflow pipeline on the CeNDR 20210121 hard-filtered isotype VCF.

Genotypes were prepared with PLINK, keeping biallelic SNVs at minor allele frequency at or above 0.05 and genotyping rate at or above 90% (--snps-only --biallelic-only --maf 0.05 --geno), and recoded to Oxford format. Both filters were computed across every strain in the phenotype file rather than across the phenotyped subset, so a marker carried by only a few of the mapped strains can still enter the scan. The Oxford genotype probabilities were converted to BIMBAM dosages as 2 x P(alternate homozygote) + P(heterozygote).

Kinship was the standardised relatedness matrix (GEMMA -gk 2), computed once per chromosome under -loco so that each chromosome’s matrix is built from the other five chromosomes only, and cached for reuse across traits. A genome-wide matrix was computed by the same call without -loco; it is not used for the association scans reported here.

Association testing was GEMMA’s univariate linear mixed model with the Wald test (-lmm 1), one call per trait per chromosome, each using that chromosome’s LOCO matrix (-k) with the marker list (-snps) and annotation (-a) built from the same PLINK output. No covariates were fitted beyond the intercept; relatedness and population structure are carried by the kinship matrix alone. Of the markers PLINK retained, the scan tested 464,045, the difference being the mitochondrial contig, which -loco excludes.

#### Initial *pos-1* RNAi experiment

A pooled population derived from sub-populations B, C, E, F, and G was thawed from a frozen stock directly into 500 ml of K media and 30 ml of 10x HB101 bacteria. After 72 hr of recovery, the gravid population was harvested and bleached in 50 ml conical tubes. Embryos were resuspended in K media and allowed to hatch and synchronize as L1s overnight. The following day, the synchronized L1s were titered and used to seed the experiment. We set up replicate liquid cultures for *pos-1* and HT115 control conditions in 250 ml baffled flasks. We put ∼75,000 worms in 50 ml of K media and 7 ml of 10X HT115 or *pos-1* RNAi bacteria in each flask and incubated them at 20°C shaking at 220 RPM for 72 hrs. Next, we bleached each of the four cultures and arrested L1s as described above. We repeated the growth conditions for the second generation. To harvest F3 L1s for sequencing, we transferred the cultures to conical tubes and allowed the F2 adults to settle for 5min. Next, we took the supernatant containing L1s and embryos through 20 µm filters, which allowed the L1s to pass and retained the embryos. We spun down the separated L1s at 1200 RPM for 2 min, removed the liquid from the cultures and transferred the pellets to microcentrifuge tubes that were stored at -80 °C until sequencing. DNA extraction, library preparation, and sequencing are described in another section.

#### Construction of an RNAi-sensitive *C. elegans* population

We used *pos-1* RNAi to evaluate *C. elegans* RNAi responses prior to pooling the strains. We thawed wild isolates that we purchased from CaeNDR on 10 cm NGMA plates seeded with OP50 and incubated them at 20°C until gravid adults were present on the plate. We bleached ∼15 adults from individual strains on 6 cm NGMA plates seeded with OP50. The following day we transferred approximately 10 L1 larvae to fresh 6 cm NGMA plates seeded with OP50 and incubated them at 20°C until L4s (F1) from the next generation were present. We then transferred 5 L4s to fresh NGMA agar plates seeded with OP50 and waited until the following generation were L4s (F2s). We transferred 10 L4 worms to NGMA + Carbomycin + IPTG plates that were seeded with induced *pos-1* RNAi bacteria. The following day we removed the adults from those plates and allowed the embryos to hatch overnight. We defined a qualitative ranking of the hatching rate we observed across the wild isolates from 0 (no embryos hatched) to 5 (all embryos hatched) and assigned each wild isolate a score. Because only 11 of the 191 tested strains had complete *pos-1* RNAi responses (score of 0), we included strains with scores of 0 or 1 to have enough strains for GWA mapping, where a score of 1 corresponds to 5 or fewer larvae hatching after *pos-1* exposure. To pool the strains we went back to the plate we sourced the worms that we placed on *pos-1* RNAi, which was starved. For each strain we decided to put in the pooled population, we chunked from the starved plates to two 10 cm NGMA plates seeded with OP50 and allowed them to grow for approximately 96 hrs until they were almost starved. We washed all the strains off of the 10 cm plates into a 50 ml conical tube and allowed the adults to settle. We aspirated the supernatant and bleached the population to harvest embryos. The following day we titered the arrested L1s and seeded a 2.5 L flask with K media, 10X HB101, and the L1 populations. We grew this population shaking at 220 RPM at 20°C for 72 hours until gravid adults were present. We bleached the entire population to harvest embryos and allowed the embryos to hatch and L1s to arrest overnight in K media. These L1s were used to seed the RNAi experiment.

#### RNAi-sensitive pooled population RNAi experiment

After we verified *pos-1* RNAi responses and harvested L1s from RNAi-sensitive individuals, we seeded flasks with the pooled population. We performed two replicates for each condition, where a replicate corresponds to a resampling of the same L1 pool. Each experiment contained 50 ml of K media, 4 ml of 10X bacteria (HT115 with an empty vector or an HT115 with gene targeting plasmid), and 75,000 L1s in a 250 ml baffled flask. We incubated the flasks in a shaking incubator at 220 RPM and at 20°C for 72 hrs. After 72 hrs we transferred the cultures to 50 ml conical tubes and allowed them to settle for 5 min, at which point we transferred 35 ml of the supernatant that contained F1 L1s. We used those L1s to seed a second generation of growth on RNAi in the same conditions described above. After another 72 hrs of growth, we separated F2 L1s from F1 adult pellets. We spun down the L1s at 1200 RPM for 2 min and saved the L1 pellet for sequencing. We also stored the F1 adult pellet for sequencing. DNA was extracted from these pellets and libraries were made as described below.

### Sequencing

Genomic DNA was prepared using Purelink Genomic DNA Mini Kit (Invitrogen cat# K1820-01), libraries were constructed using the Nextera XT library kit (illumina cat# FC-131-1024), and sequenced on the NextSeq2000 platform.

### Cross Construction

Before cross-construction, male stocks were generated by heat-shock-induced nondisjunction. For each strain, fifteen L4 hermaphrodites were transferred to three 6 cm NGMA plates and incubated at 30°C for 5 hours, and then returned to 20°C. After 72 hours, plates were screened for males and crossed to L4 hermaphrodites maintained at 20°C. For each male cross, 5-10 hermaphrodites and 10-20 males were placed on 6 cm NGMA plates that were seeded with 10 µL of OP50 and allowed to mate overnight. The following day, individual mated hermaphrodites were transferred to fresh 6 cm NGMA plates, and the resulting progeny were monitored for an approximately 50:50 male-to-hermaphrodite ratio. Crosses for population construction were performed using the same mating procedure described above.

JU2466 was the parent JU1793 x JU2466 and N2 x XZ1516. Each strain serves once as the hermaphrodite parent and once as the male parent. After mating, an equal number of plugged hermaphrodites were pooled within each reciprocal pair and plated onto 10 cm NGMA plates, yielding two independently amplified cross populations.

### Cross Maintenance

Crosses were propagated for nine generations on 10 cm NGMA plates seeded with 1mL of OP50, with approximately 100,000 worms maintained per generation. When abundant L1 larvae were present, plates were washed with M9 buffer and gravity-settled to separate higher-density, older worms in the pellet from lower-density L1 larvae in the supernatant that would be seeded for the next generation. L1 larvae were titered prior to plating (∼1,000 L1s per NGMA plate), and remaining larvae were intermittently frozen down for long-term storage.

For generation nine, the number of plated worms was doubled to ensure sufficient population size for three RNAi conditions and a time-zero sequencing control. Ten thousand L1s from the resulting F10 progeny were distributed across the three RNAi conditions and propagated using the same procedure, with remaining larvae from each generation saved for downstream sequencing.

### NIL Construction

To validate the QTL on Chromosome III, multiple sets of Near-Isogenic Lines (NILs) were constructed and phenotyped. Heterozygous F1 animals were first generated by crossing JU1793 and JU2466. Approximately 72 hours later, F2 recombinant progeny were isolated by transferring single F2 animals into separate wells of a 96-well plate containing 50 uL of K media and 0.008 mg/ml HB101. After each well produced F3 progeny and reached starvation, 15 uL of the worm suspension was added to 15 uL of 2x lysis buffer containing 2% proteinase K. Samples were processed in a thermocycler at 60°C for 1 hour, followed by heat inactivation at 95°C for 20 minutes. After genotyping with primers targeting the regions of interest (Table S25 - strains; Table S26 - oligos), starved F3 suspensions were recovered onto 6 cm NGMA plates seeded with OP50. Recovered F3 progeny were singled and re-genotyped to identify homozygous recombinant individuals. Confirmed NIL candidates were then backcrossed to their respective parental strain to reduce background variation further. Once the introgressed interval became too small for additional recombination, CRISPR-Cas9-mediated homologous recombination was used for further fine-mapping [40]. Guide RNAs and PCR oligos to check for recombinant progeny are outlined in Table S26.

### CRISPR Design to edit *sid-2*

We used the guide RNA gSZ182 to create a double stranded break near the position of the III:13680248 variant that differentiates JU1793 and JU2466 [JU1793:C>JU2466:A] at the SID-2 locus. Injection mixes were made with repair templates oZ446 or oZ456 to swap the JU1793 allele to the JU2466 allele and the JU2466 allele to the JU1793 allele, respectively. Injected animals were singled 16 hours after injection and allowed to generate progeny. Roller F1 progeny were singled and allowed to lay progeny for 24 hrs at which point the F1 roller parent was lysed for DNA extraction. The primers oZ461 and oZ468 were used to amplify the region surrounding the edit site. PCR amplicons (10 µl) were subsequently digested with HpyCH4IV using the cutsmart buffer.

### Plate-based *pos-1* embryonic lethality assay

Strains were synchronized by hypochlorite treatment and grown until L4s were visible. Three days prior to plating onto RNAi bacteria, populations were amplified by transferring five L4s to each of three plates. After approximately 72 hours, 10-15 L4s were plated onto each RNAi condition. The following day, 150 and 200 embryos were transferred to an unseeded NGMA plate and counted to determine the exact starting number of embryos. After 24 hours, the number of unhatched embryos was counted. RNAi sensitivity was quantified as the fraction of embryos remaining unhatched divided by the total number of embryos plated.

## Author Contributions

**Conceptualization:** Stefan Zdraljevic; Joshua S. Bloom

**Data curation:** Stefan Zdraljevic

**Formal analysis:** Stefan Zdraljevic; Joshua S. Bloom

**Investigation:** Stefan Zdraljevic; Alex Lee; Laura Walter-McNeill; Heriberto Marquez

**Software:** Stefan Zdraljevic; Joshua S. Bloom

**Visualization:** Stefan Zdraljevic

**Methodology:** Stefan Zdraljevic; Joshua S. Bloom

**Writing – original draft:** Stefan Zdraljevic

**Writing – review and editing:** Stefan Zdraljevic; Joshua S. Bloom; Leonid Kruglyak

**Project administration:** Stefan Zdraljevic; Joshua S. Bloom; Leonid Kruglyak

**Supervision:** Stefan Zdraljevic; Joshua S. Bloom; Leonid Kruglyak

**Resources:** Stefan Zdraljevic; Amy K. Webster; Rojin Chitrakar; Jameson Blount; L. Ryan Baugh; Leonid Kruglyak

**Funding acquisition:** Stefan Zdraljevic

## Data and materials availability

All data are available in the manuscript or the supplementary materials.

---

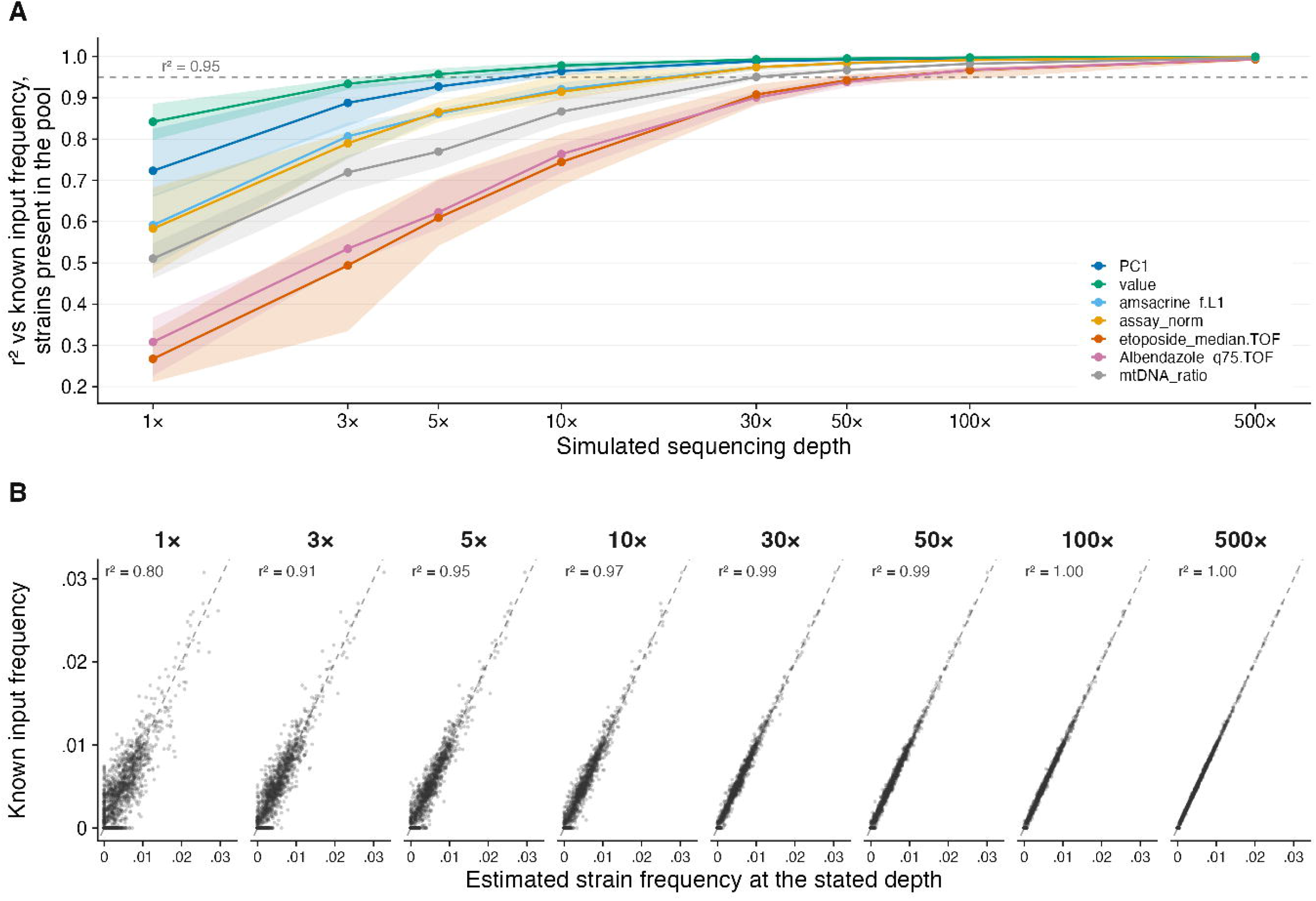

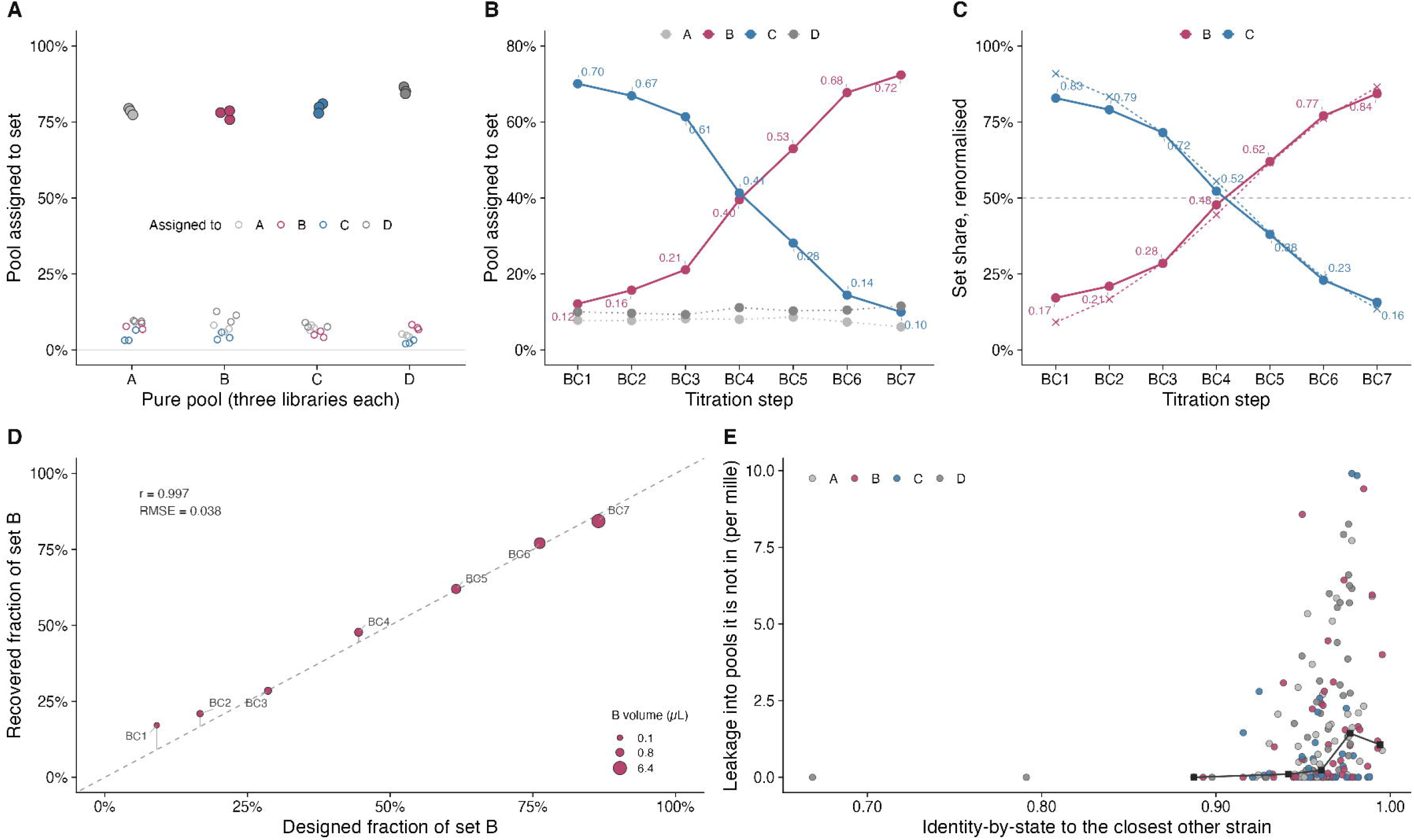

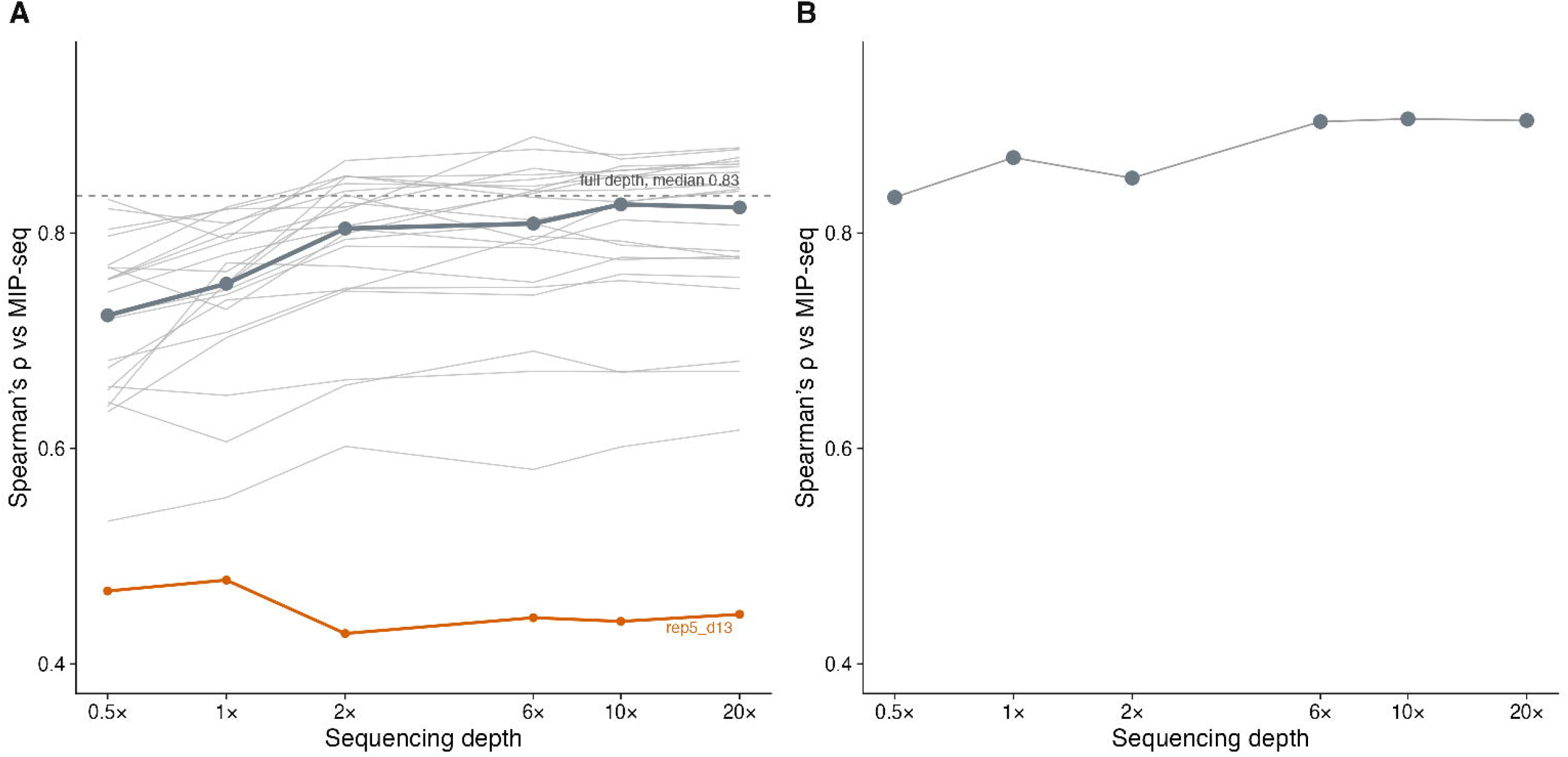

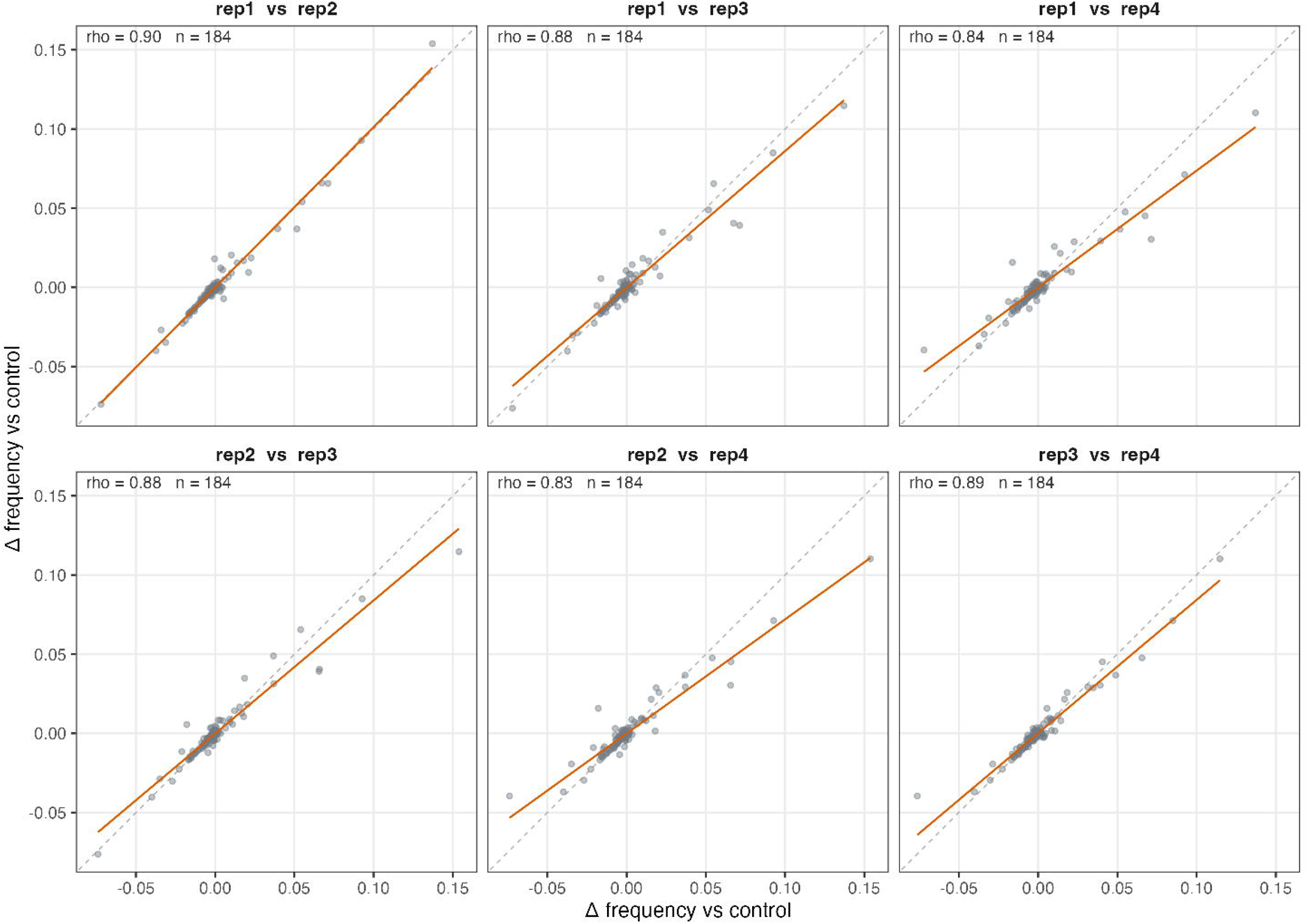

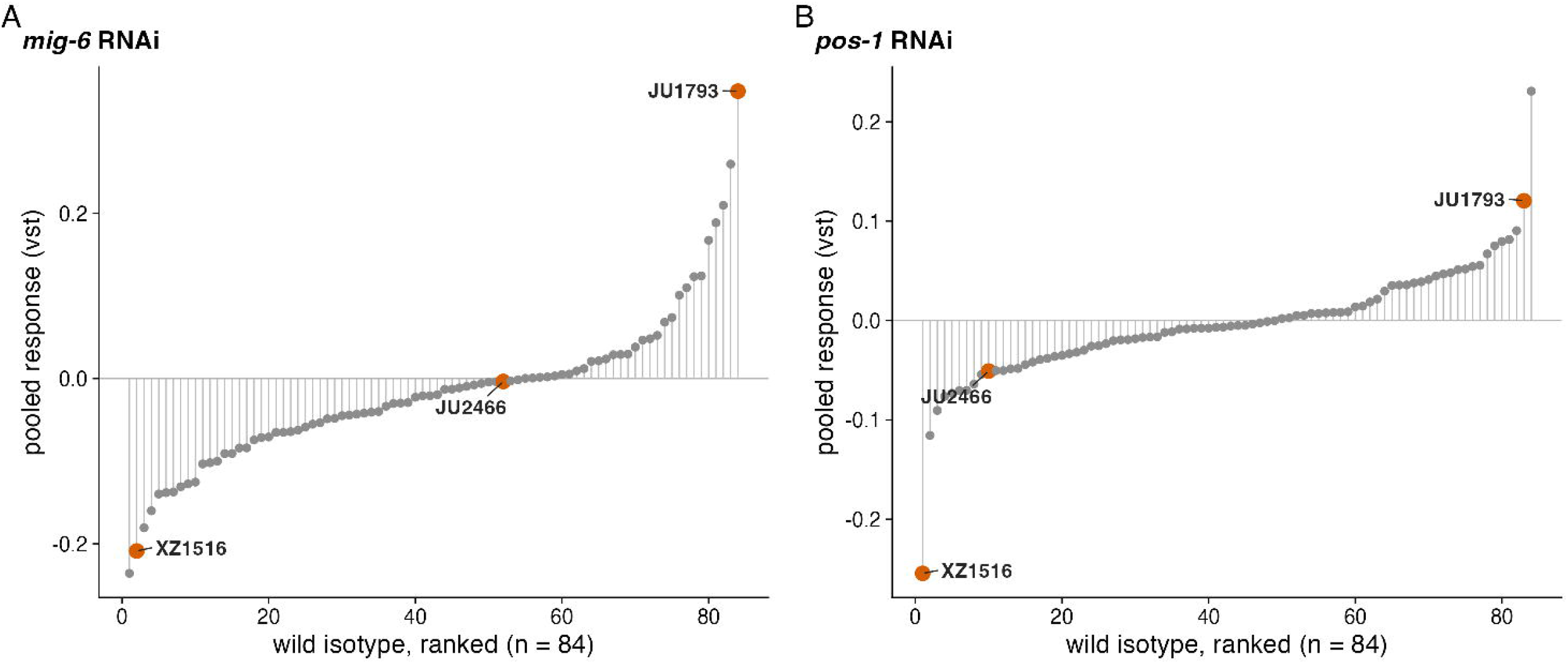

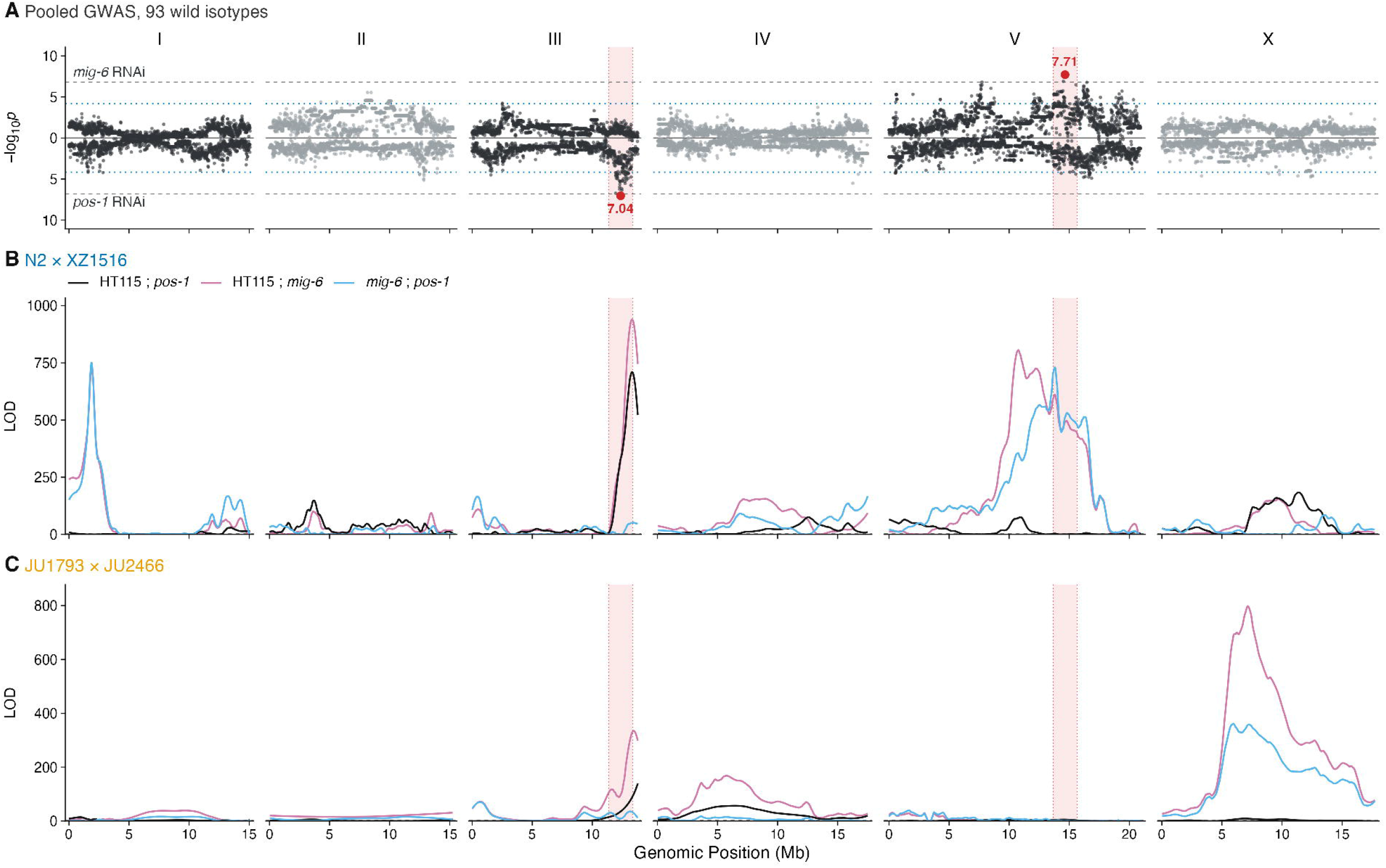

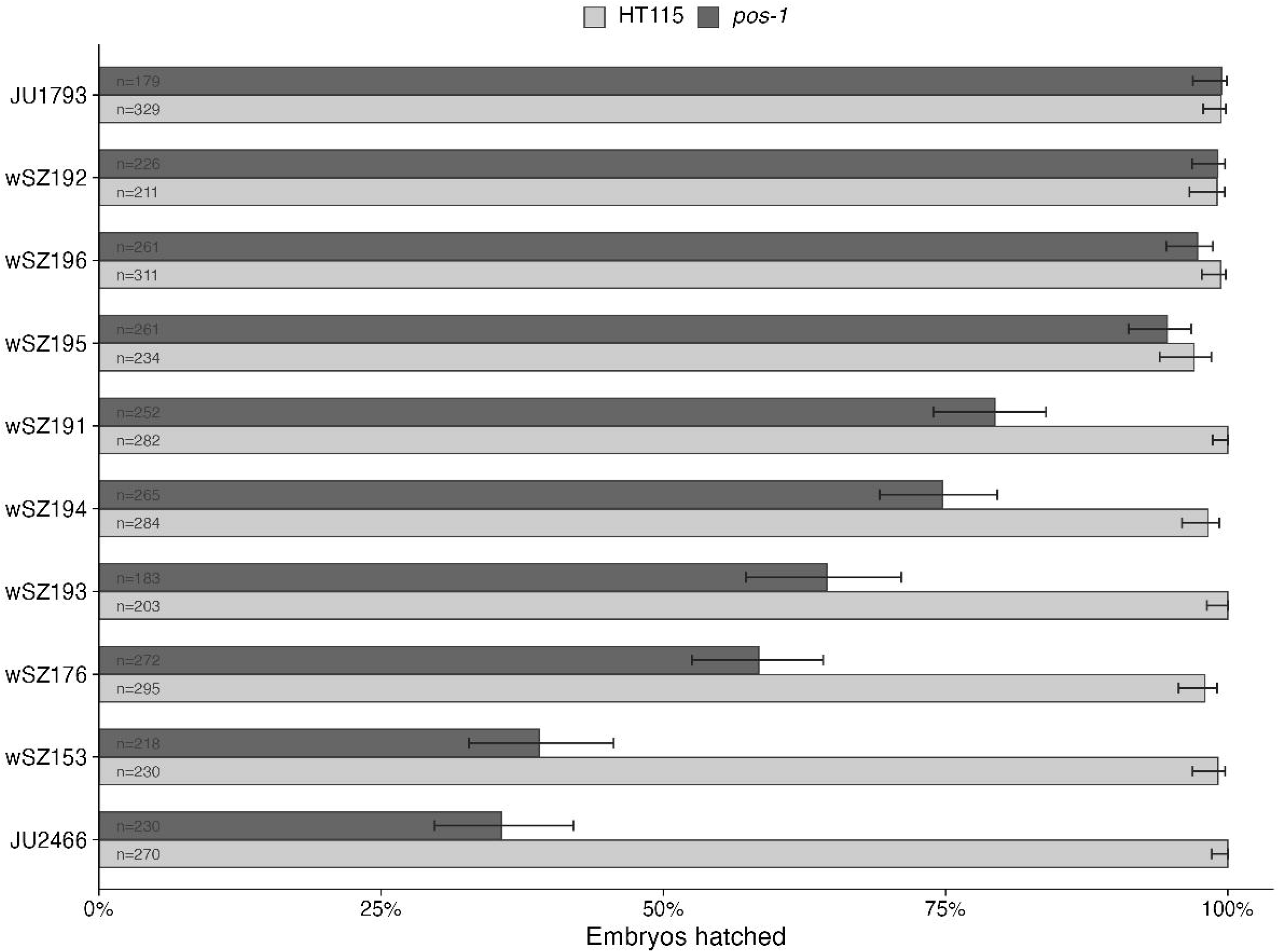

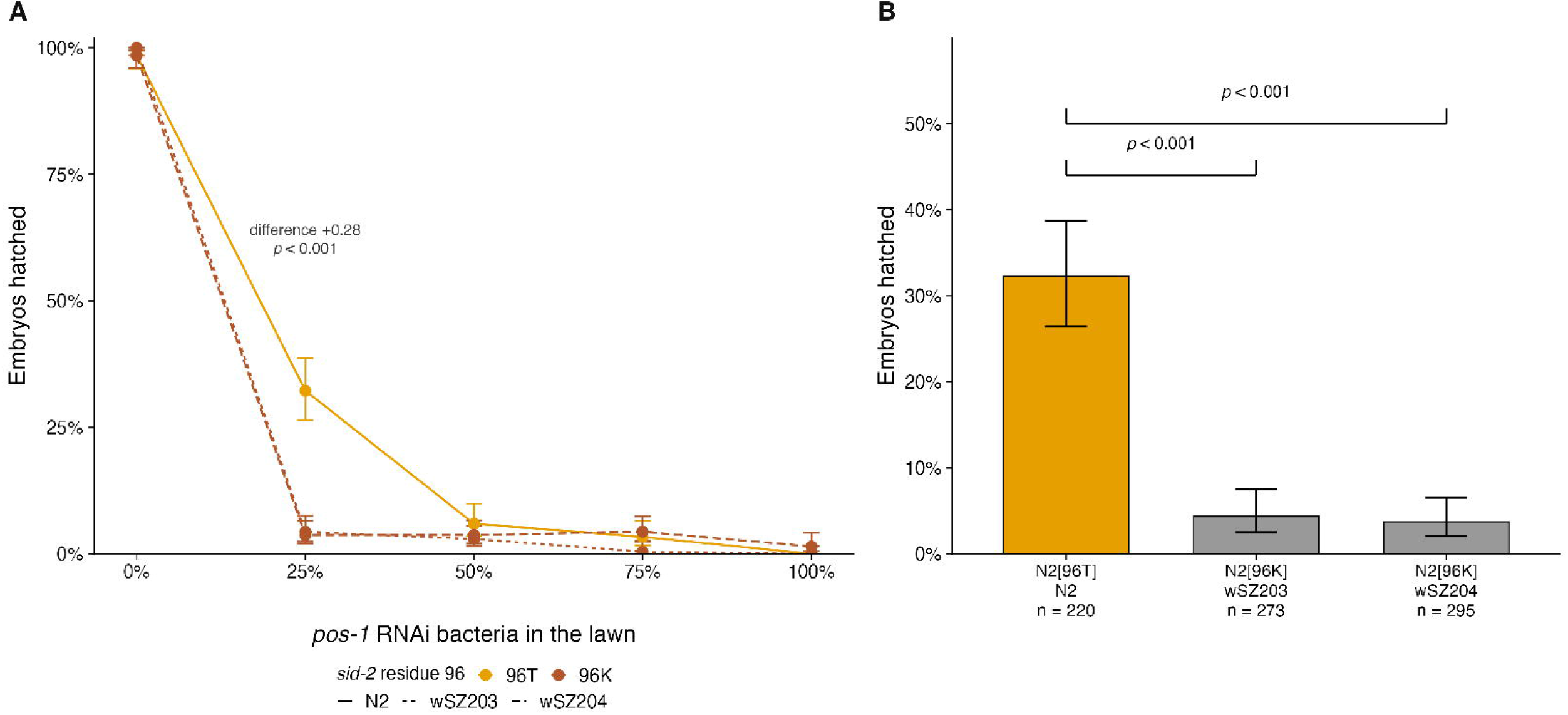

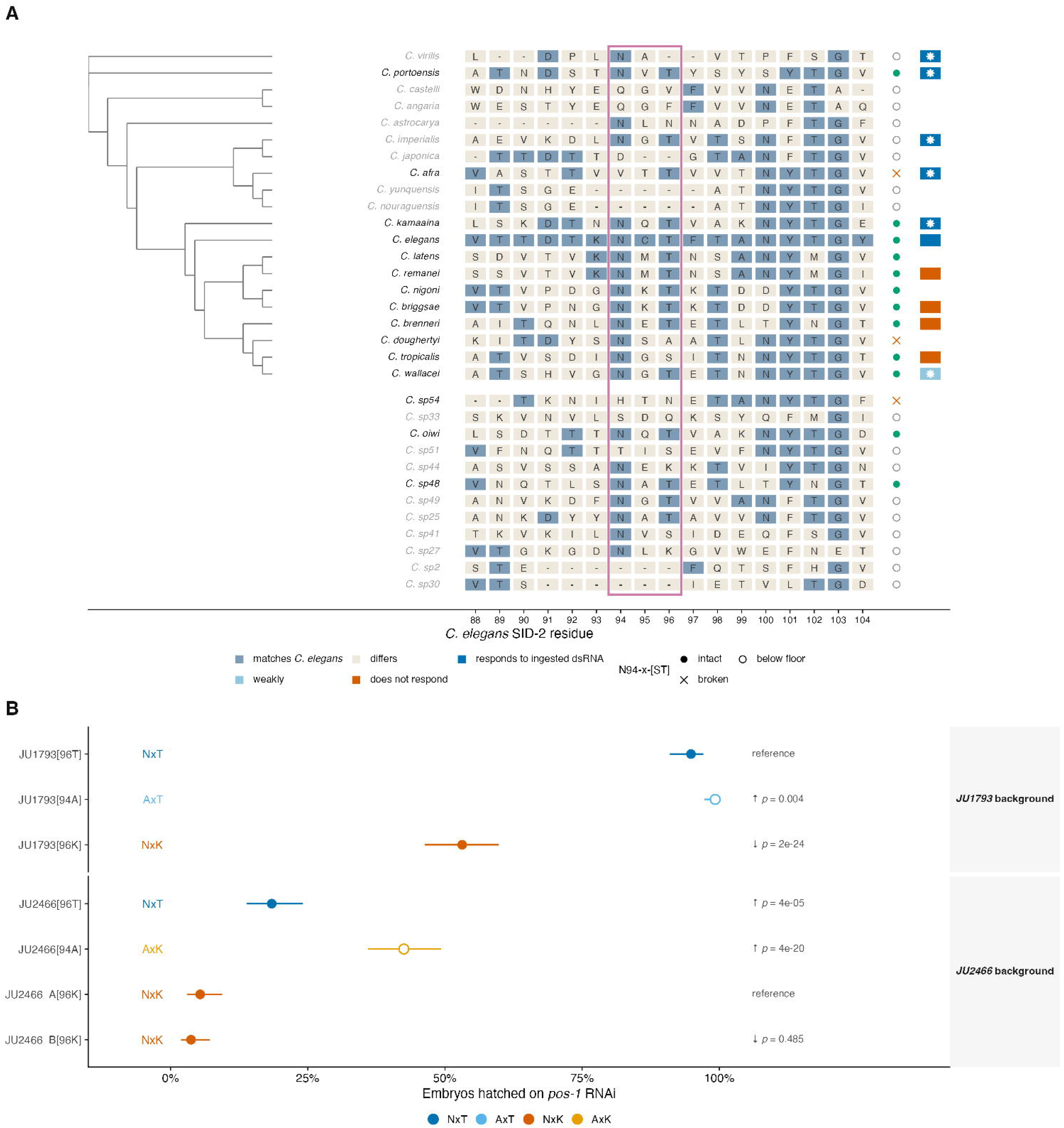

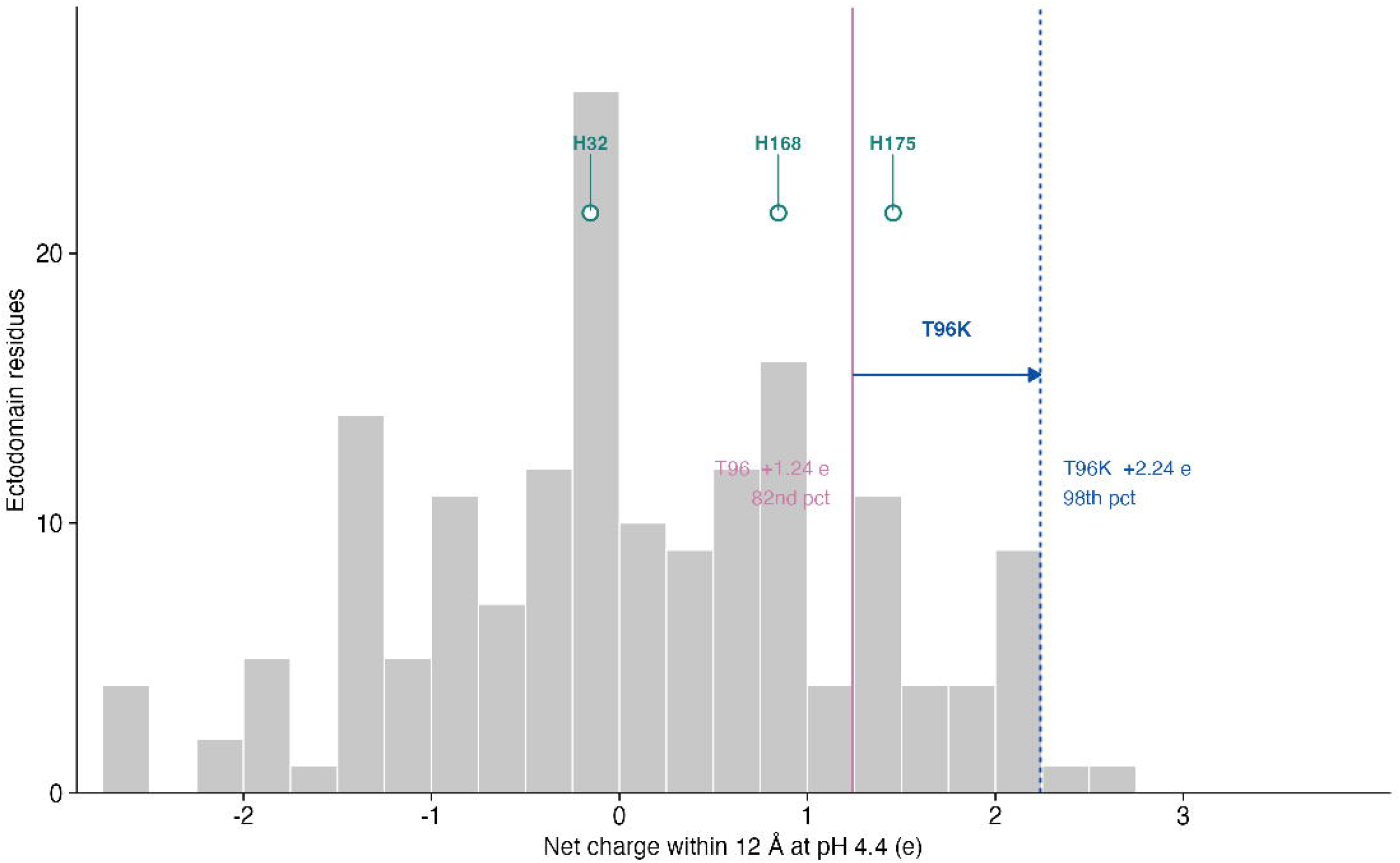

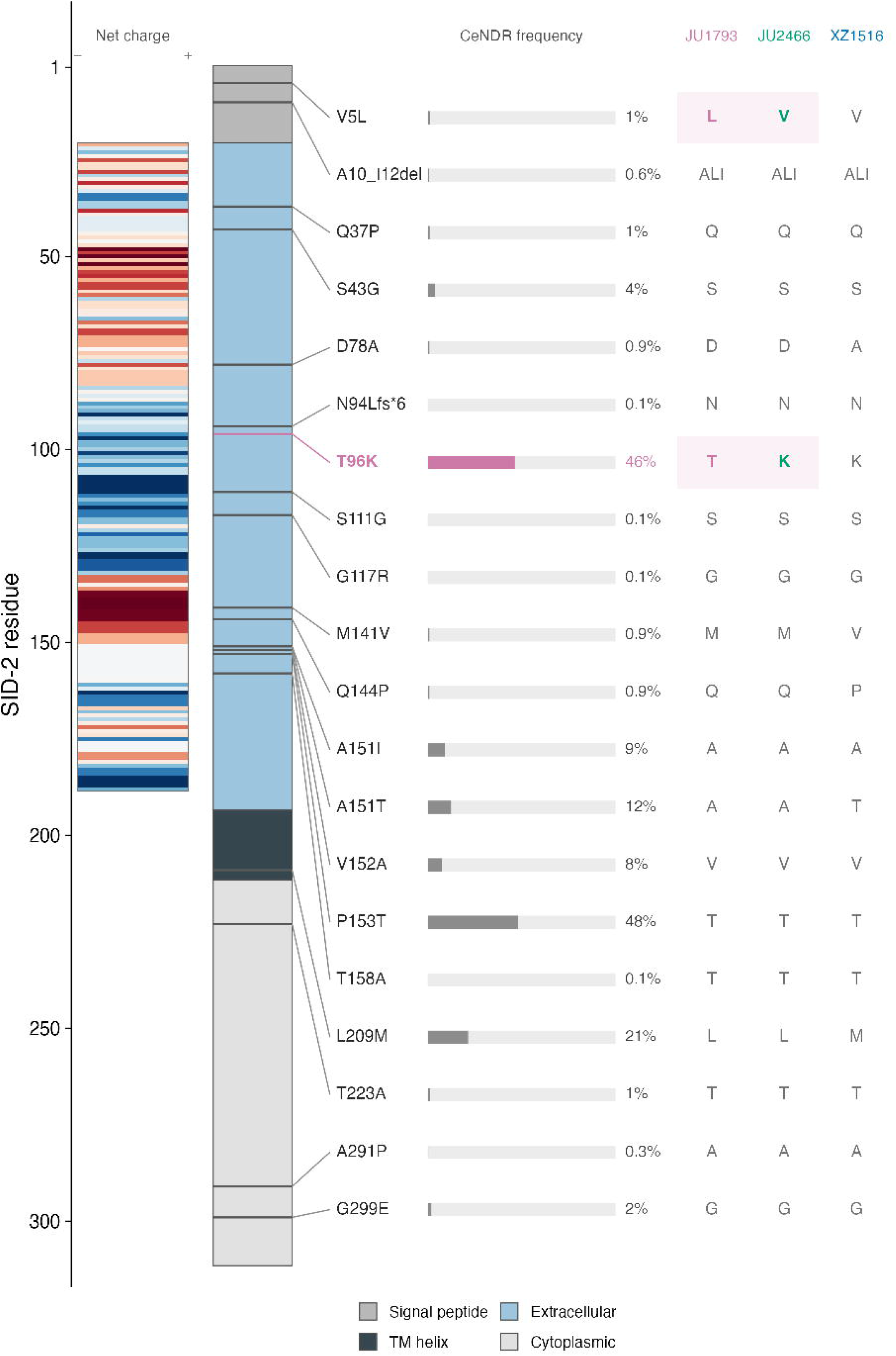

