## Supplementary material for "Large-scale pooled selection enables mapping of modifiers of RNAi gene knockdown effects in *C. elegans*": README.md

# Manuscript supplement

Supplemental tables for *Flexible pooled phenotyping enables population-scale
mapping of RNAi-sensitivity modifiers*, numbered in order of first mention in
the Results.

Built by `scripts/make_manuscript_supplement.py` from `supplemental_data/` and
`plots/`. Rerunning it rebuilds this directory from scratch, so edit the script
rather than the files here. The script reads a flattened text extraction of
`Bulk Paper.pdf` at `/tmp/bulk_flat.txt` in order to check every anchor against
the manuscript; regenerate it with:

```sh
python3 -c "
import re, pypdf
r = pypdf.PdfReader('Bulk Paper.pdf')
t = '\n'.join((p.extract_text() or '') for p in r.pages)
open('/tmp/bulk_flat.txt','w').write(re.sub(r'\s+',' ',t))"
```

**Cite near** gives a short verbatim phrase from the manuscript, checked against
the text at build time. Paste it into a find box to land on the sentence where
the table reference belongs.

`TABLE_INDEX.csv` is the same list as a spreadsheet.

`kind` separates the two things in here. A **table** is small and opens in a
spreadsheet, and is what a journal expects as a supplemental table. A
**dataset** is either large or in an R binary format; those belong in the data
repository the paper deposits to, and should be cited under data availability
rather than attached to the article. The datasets are most of the bulk here.

### Table S1 — `Table_S01_pos1_pilot_strain_phenotypes.csv.gz`

Per-strain pos-1 RNAi response for the 231-isotype pilot pool. Columns: delta_ctrl_pos-1_T2 (RNAi minus control change in pool frequency), vst_ctrl_pos-1_T2 (variance-stabilised, the trait the association scan was run on), log2fc_ctrl_pos-1_T2 (the log2 ratio), and two negative-control columns, growth on HT115 against t0 and log10 control abundance.

**Cite near:** “231 pooled wild isolates”

*Source in repository:* `supplemental_data/phenotypes/pos1_2023_association_traits.csv.gz`  (0.01 MB)

### Table S2 — `Table_S02_simulation_recovery_by_depth_a.tsv; Table_S02_simulation_recovery_by_depth_b.tsv`

Deconvolution recovery in simulation. r-squared of estimated against known input frequency for each of seven fitness traits at eight sequencing depths, over ten seeded replicates, plus the seven trait vectors used as fitness and the strains carrying each.

**Cite near:** “NNLS recovered strain frequencies”

*Source in repository:* `supplemental_data/deconvolution/simulation_seeded_r2.tsv; supplemental_data/deconvolution/simulation_fitness_traits.tsv`  (0.05 MB)

### Table S3 — `Table_S03_known_mixture_design_and_recovery_a.tsv; Table_S03_known_mixture_design_and_recovery_b.tsv; Table_S03_known_mixture_design_and_recovery_c.tsv.gz`

The designed DNA mixture, in three parts: (a) the titration design, giving each sample's set B and set C volumes, water, total DNA mass and nominal set fractions; (b) which of the four sets each strain belongs to, with its isotype; (c) the recovered per-strain frequency in every library.

**Cite near:** “root mean squared error”

*Source in repository:* `supplemental_data/deconvolution/dilution_design.tsv; supplemental_data/deconvolution/dilution_strain_sets.tsv; supplemental_data/deconvolution/dilution_predictions_poolref.tsv.gz`  (0.03 MB)

### Table S4 — `Table_S04_nnls_vs_mipseq_frequencies_a.tsv.gz; Table_S04_nnls_vs_mipseq_frequencies_b.txt.gz`

Per-strain, per-sample frequencies from this platform (NNLS on whole-genome sequence) beside the published MIP-seq frequencies for the same 23 samples of the L1 starvation time course. The basis for the platform comparison.

**Cite near:** “highly correlated with those derived from MIP-seq”

*Source in repository:* `supplemental_data/deconvolution/baugh_nnls_dep103_with_mipseq.tsv.gz; supplemental_data/deconvolution/mipseq_frequencies.txt.gz`  (0.08 MB)

### Table S5 — `Table_S05_downsampling_recovery.tsv`

Recovery of the two published traits at each subsampled depth (0.5, 1, 2, 6, 10 and 20x): Spearman correlation against the published PC1 and Slope, and against Slope computed on the difference-based definition used here.

**Cite near:** “as we downsampled reads”

*Source in repository:* `supplemental_data/deconvolution/baugh_downsample_trait_recovery.tsv`  (0.00 MB)

### Table S6 — `Table_S06_pos1_pilot_replicate_frequencies.csv.gz`

Sample-level strain frequencies for the pilot pool: four pos-1 replicate pools and two control pools at three read-depth cutoffs. Supports the replicate-agreement comparison and the per-isolate responsiveness counts.

**Cite near:** “across all six pairwise replicate comparisons”

*Source in repository:* `supplemental_data/phenotypes/pos1_2023_sample_frequencies.csv.gz`  (0.15 MB)

### Table S7 — `Table_S07_pos1_pilot_gwas.csv.gz`  *(dataset -- deposit, not an article supplement)*

Genome-wide association results for the pilot pos-1 response, 464,045 markers over 231 strains, GEMMA with leave-one-chromosome-out kinship.

**Cite near:** “genome-wide association scan on the pos-1 responses”

*Source in repository:* `supplemental_data/mapping/pos1_2023_gemma_loco.csv.gz`  (7.14 MB)

### Table S8 — `Table_S08_gwas_significance_thresholds.tsv`

Effective number of independent tests by the Li and Ji (2005) eigenvalue method, per panel and per chromosome, with the marker counts, the imputed fraction, the trace check, and the resulting Bonferroni and eigenvalue thresholds drawn on every association panel.

**Cite near:** “passed the Bonferroni significance threshold”

*Source in repository:* `supplemental_data/mapping/eigen_independent_tests.tsv`  (0.00 MB)

### Table S9 — `Table_S09_manual_plate_scores.tsv`

Manual pos-1 RNAi score per wild isolate, on a six-level ordinal scale where 0 is a complete RNAi response and 5 is none. Two columns, strain and score; which strains were carried into the 93-strain pool is not recorded here.

**Cite near:** “191 wild strains”

*Source in repository:* `supplemental_data/phenotypes/plate_scores_pos1.tsv`  (0.00 MB)

### Table S10 — `Table_S10_paaby2015_comparison.txt.gz`

Well-level embryo and larva counts from Paaby et al. 2015, restricted to the pos-1 clone, as used for the independent comparison against the manual plate scores.

**Cite near:** “previously published evaluation of wild isolate RNAi responses”

*Source in repository:* `supplemental_data/phenotypes/paaby2015_embryonic_lethality.txt.gz`  (0.58 MB)

### Table S11 — `Table_S11_pooled_ten_target_phenotypes.csv.gz`

Per-strain responses for the 93-strain pool across all ten RNAi targets against the HT115 empty-vector control, one column per target in each of the three parameterisations (delta_ctrl, log2fc, vst). The vst columns are what the association scans were run on.

**Cite near:** “each of ten target genes”

*Source in repository:* `supplemental_data/phenotypes/pooled_vst_traits.csv.gz`  (0.03 MB)

### Table S12 — `Table_S12_pooled_gwas_and_cross_scans.rds`  *(dataset -- deposit, not an article supplement)*

The thinned mapping bundle behind Figure 2: pooled association scans for all ten targets and the cross contrast scans, one marker per bin retained on maximum LOD so every peak survives. Peak positions and heights are identical to the full data; interval widths may be 0-9 kb narrower.

**Cite near:** “exceeded the Bonferroni threshold for pos-1 and mig-6”

*Source in repository:* `supplemental_data/mapping/pooled_cross_bundle_thinned.rds`  (9.64 MB)

### Table S13 — `Table_S13_cross_allele_frequencies_a.tsv.gz; Table_S13_cross_allele_frequencies_b.tsv.gz; Table_S13_cross_allele_frequencies_c.tsv`  *(dataset -- deposit, not an article supplement)*

Genome-wide parental allele frequencies for both advanced-intercross crosses, per sample, with the sample sheet naming the condition and timepoint of each.

**Cite near:** “allele frequency deviations between conditions”

*Source in repository:* `supplemental_data/mapping/cross_af_JU1793xJU2466.tsv.gz; supplemental_data/mapping/cross_af_N2xXZ1516.tsv.gz; supplemental_data/mapping/cross_af_samples.tsv`  (11.71 MB)

### Table S14 — `Table_S14_cross_qtl_summary.tsv`

THE TABLE THE TEXT ALREADY CITES AS "Supplemental table X". Every cross QTL peak above the genome-wide threshold in every contrast (495 rows): position, peak LOD, support interval, parental frequencies either side, rank on its chromosome, whether the trough separates it from a taller peak, and whether the other cross calls it by interval overlap or by position.

**Cite near:** “Supplemental table X”

*Source in repository:* `plots/TABLE_cross_qtl_full.tsv`  (0.13 MB)

### Table S15 — `Table_S15_cross_locus_classification_a.tsv; Table_S15_cross_locus_classification_b.tsv`

General against target-specific classification of each cross locus: how many RNAi targets respond at it and in which direction, with the per-condition frequency differences the call is made on, swept over four response cutoffs.

**Cite near:** “QTL that are shared between conditions”

*Source in repository:* `plots/TABLE_cross_qtl_locus_classification.tsv; plots/TABLE_cross_qtl_condition_dfreq.tsv`  (0.09 MB)

### Table S16 — `Table_S16_nil_introgression_ranges.bed`

Introgression boundaries for the NIL series as BED: chromosome, start, end, strain and donor parent, with the two parents carried as whole-chromosome rows. These are the sequence-confirmed lines, and the difference between the wSZ191 and wSZ196 rows is the 37 kb interval. Which boundaries are sequence-confirmed as against marker-bounded is in Table S25, not here.

**Cite near:** “37 kb interval”

*Source in repository:* `supplemental_data/hatching_assays/nil_introgression_ranges.bed`  (0.00 MB)

### Table S17 — `Table_S17_nil_hatching_counts.tsv`

Embryo hatching for every NIL and both parents on pos-1 RNAi and on control food. Columns: strain, condition, embryos plated, number unhatched, and hatched fraction. One plate per strain per condition, so the binomial intervals quoted in the text are computed from these counts rather than stored here.

**Cite near:** “distinct pos-1 RNAi phenotypes”

*Source in repository:* `supplemental_data/hatching_assays/nil_series_hatching.tsv`  (0.00 MB)

### Table S18 — `Table_S18_nil_interval_content_a.tsv; Table_S18_nil_interval_content_b.tsv; Table_S18_nil_interval_content_c.tsv`

What the 37 kb interval contains: every gene, every difference between the cross parents with its annotated consequence and impact class, and the exon models. The basis for the two protein-altering differences in sid-2.

**Cite near:** “27 parental differences across twelve genes”

*Source in repository:* `supplemental_data/mapping/nil_interval_genes.tsv; supplemental_data/mapping/nil_interval_parent_variants.tsv; supplemental_data/mapping/nil_interval_exons.tsv`  (0.01 MB)

### Table S19 — `Table_S19_sid2_allele_swap_hatching_ju.csv`

Embryo hatching for the JU1793 and JU2466 sid-2 allele swaps and their parents on pos-1 and control food. Columns: experiment, strain, genotype, glycosylation motif, condition, embryos plated, number unhatched, hatched fraction. The motif column is what distinguishes the residue-94 and residue-96 states (NxT, NxK, AxT, AxK) and records that JU2466 appears as two isolates, A and B, with the 96T edit made in A.

**Cite near:** “reciprocal allele-swap strains”

*Source in repository:* `supplemental_data/hatching_assays/ju_allele_swaps_hatching.csv`  (0.00 MB)

### Table S20 — `Table_S20_sid2_allele_swap_hatching_n2.tsv`

Embryo hatching for N2 and the two N2 sid-2 96K lines (wSZ203, wSZ204) across the 0, 25, 50, 75 and 100% pos-1 food dose series. One row per strain per dose; the pooled 96K figure quoted in the text is computed from the two lines' counts rather than stored as a row.

**Cite near:** “lowered the N2 hatching rate”

*Source in repository:* `supplemental_data/hatching_assays/n2_allele_swaps_hatching.tsv`  (0.00 MB)

### Table S21 — `Table_S21_sid2_ortholog_conservation_a.tsv; Table_S21_sid2_ortholog_conservation_b.tsv; Table_S21_sid2_ortholog_conservation_c.tsv; Table_S21_sid2_ortholog_conservation_d.tsv`

Conservation of the N94-C95-T96 sequon across the Caenorhabditis species surveyed, with the ortholog search results, the alignable-window survey bounding how far SID-2 can be compared, and the species name map.

**Cite near:** “Elegans supergroup”

*Source in repository:* `supplemental_data/structure/sid2_ortholog_conservation.tsv; supplemental_data/structure/sid2_ortholog_search.tsv; supplemental_data/structure/sid2_ortholog_window_survey.tsv; supplemental_data/structure/sid2_species_name_map.tsv`  (0.02 MB)

### Table S22 — `Table_S22_sid2_environmental_competence.tsv`

Published environmental-RNAi competence per Caenorhabditis species, curated from the literature: the species, the strain tested for RNAi response, the strain whose genome was sequenced, whether those are the same isolate, the reported response, the kind of evidence, and the source. Layered onto the conservation figure.

**Cite near:** “sensitive to ingested double stranded RNA”

*Source in repository:* `supplemental_data/structure/sid2_env_rnai_sensitivity.tsv`  (0.00 MB)

### Table S23 — `Table_S23_sid2_local_charge_a.tsv; Table_S23_sid2_local_charge_b.tsv`

Local net charge in 12 A windows across the SID-2 ectodomain, per residue, at two pH values (4.4 and 7.4), with the number of residues inside each window, the model pLDDT and the residue coordinates. The second file carries the per-residue model quantities behind it: pLDDT, assigned secondary structure, topology region and coordinates.

**Cite near:** “local net charge in 12”

*Source in repository:* `supplemental_data/structure/sid2_local_charge.tsv; supplemental_data/structure/sid2_per_residue.tsv`  (0.03 MB)

### Table S24 — `Table_S24_sid2_population_variants_a.csv; Table_S24_sid2_population_variants_b.tsv; Table_S24_sid2_population_variants_c.tsv; Table_S24_sid2_population_variants_d.tsv`

sid-2 variation across the wild population from CaeNDR. (a) The CaeNDR 20250625 variant export for sid-2, verbatim: every variant in the gene, its consequence, and the isotypes carrying it, the source of Figure 4D and Figure S12 (frequencies are carriers over the release's 684 isotypes). (b-d) The earlier annotation: protein-altering variants segregating among the cross parents with 20210121 frequencies, the population missense set from 20231213, and the differences between the parents of each cross.

**Cite near:** “high-frequency variant in the dsRNA transporter SID-2”

*Source in repository:* `supplemental_data/structure/sid2_cendr20250625_variants.csv; supplemental_data/structure/sid2_variants_cendr.tsv; supplemental_data/structure/sid2_population_missense.tsv; supplemental_data/structure/sid2_parental_variants.tsv`  (0.04 MB)

### Table S25 — `Table_S25_strains.csv`

Every strain constructed for this work: QX designation, lab identifier, introgression or allele designation, genotype, background, construction route, guide and repair template used, and verification status.

**Cite near:** “Methods: NIL Construction”

*Source in repository:* `supplemental_data/Table_S25_strains.csv`  (0.01 MB)

### Table S26 — `Table_S26_oligonucleotides.csv`

Every oligonucleotide: genotyping primers with their pairings and expected products, restriction assays, the four sid-2 repair templates and the four guide RNAs.

**Cite near:** “CRISPR Design to edit sid-2”

*Source in repository:* `supplemental_data/Table_S26_oligonucleotides.csv`  (0.00 MB)


## Not promoted

60 files in `supplemental_data/` support no manuscript figure or claim and stay where they are -- alternative deconvolution references, the mig-6 locus census that no cited figure draws, intermediate caches, and the structure files behind figures the manuscript does not cite.

- `supplemental_data/SUPPLEMENTAL_DATA_OVERVIEW.md`
- `supplemental_data/Strains_Oligos.csv`
- `supplemental_data/deconvolution/baugh_bootstrap_array.rda`
- `supplemental_data/deconvolution/baugh_downsampled_slopes.rda`
- `supplemental_data/deconvolution/baugh_nnls_dep102_with_mipseq.tsv.gz`
- `supplemental_data/deconvolution/baugh_nnls_pool100_with_mipseq.tsv.gz`
- `supplemental_data/deconvolution/baugh_nnls_with_mipseq.RData`
- `supplemental_data/deconvolution/baugh_strain_order.txt`
- `supplemental_data/deconvolution/baugh_strain_private_markers.tsv`
- `supplemental_data/deconvolution/baugh_strain_similarity.tsv`
- `supplemental_data/deconvolution/cache_boot_freq.rds`
- `supplemental_data/deconvolution/cache_boot_slopes.rds`
- `supplemental_data/deconvolution/dilution_predictions_bcref.tsv.gz`
- `supplemental_data/deconvolution/dilution_predictions_fullref.tsv.gz`
- `supplemental_data/deconvolution/dilution_predictions_regenotype.tsv.gz`
- `supplemental_data/deconvolution/dilution_strain_similarity.tsv`
- `supplemental_data/deconvolution/simulation_gwas_traits.tsv.gz`
- `supplemental_data/deconvolution/simulation_nnls_frequencies.tsv.gz`
- `supplemental_data/deconvolution/simulation_reported_r2.tsv`
- `supplemental_data/deconvolution/simulation_seeded_frequencies.tsv.gz`
- `supplemental_data/genotypes/gwas_peak_genotypes.tsv`
- `supplemental_data/genotypes/sid2_region.bed`
- `supplemental_data/genotypes/sid2_region.bim`
- `supplemental_data/genotypes/sid2_region.fam`
- `supplemental_data/genotypes/sid2_region.log`
- `supplemental_data/mapping/ju_cross_ht115_vs_pos1_scan.tsv.gz`
- `supplemental_data/mapping/jx_cross_chr3_peaks.tsv`
- `supplemental_data/mapping/jx_cross_chr3_profile.tsv.gz`
- `supplemental_data/mapping/jx_cross_sid2_window.tsv`
- `supplemental_data/mapping/mig6_locus_divergent.tsv`
- `supplemental_data/mapping/mig6_locus_exons.tsv`
- `supplemental_data/mapping/mig6_locus_genes.tsv`
- `supplemental_data/mapping/mig6_locus_nocall.tsv`
- `supplemental_data/mapping/mig6_locus_summary.tsv`
- `supplemental_data/mapping/mig6_locus_variants.tsv`
- `supplemental_data/phenotypes/baugh_association_traits.csv`
- `supplemental_data/phenotypes/baugh_association_traits_dep103.csv`
- `supplemental_data/phenotypes/baugh_association_traits_pool100.csv`
- `supplemental_data/phenotypes/baugh_mapping_traits.csv`
- `supplemental_data/phenotypes/baugh_published_traits.txt`
- `supplemental_data/phenotypes/baugh_recipe_traits_dep103.csv`
- `supplemental_data/phenotypes/baugh_recipe_traits_pool100.csv`
- `supplemental_data/reagent_table_issues.txt`
- `supplemental_data/structure/sid2_alphafold_confidences/fold_2026_02_17_15_52_summary_confidences_0.json`
- `supplemental_data/structure/sid2_alphafold_confidences/fold_2026_02_17_15_52_summary_confidences_1.json`
- `supplemental_data/structure/sid2_alphafold_confidences/fold_2026_02_17_15_52_summary_confidences_2.json`
- `supplemental_data/structure/sid2_alphafold_confidences/fold_2026_02_17_15_52_summary_confidences_3.json`
- `supplemental_data/structure/sid2_alphafold_confidences/fold_2026_02_17_15_52_summary_confidences_4.json`
- `supplemental_data/structure/sid2_alphafold_dimer.pdb`
- `supplemental_data/structure/sid2_deeptmhmm_topology.3line`
- `supplemental_data/structure/sid2_membrane_oriented.pdb`
- `supplemental_data/structure/sid2_ortholog_alignment.tsv`
- `supplemental_data/structure/sid2_ortholog_sequences.fa`
- `supplemental_data/structure/sid2_panelC_published.tsv`
- `supplemental_data/structure/sid2_panelC_residues.tsv`
- `supplemental_data/structure/sid2_sequences.fa`
- `supplemental_data/structure/sid2_species_alignment.tsv`
- `supplemental_data/structure/sid2_species_tree.nwk`
- `supplemental_data/structure/sid2_variant_ld.tsv`
- `supplemental_data/structure/sid2_variants_cendr_README.txt`
